# Chromosome scale genomes of two invasive Adelges species enable virtual screening for selective adelgicides

**DOI:** 10.1101/2024.11.21.624573

**Authors:** AM Glendening, C Stephens, VS Vuruputoor, T Chaganti, MN Myles, DL Stern, M Abdelalim, Y-P Juang, SA Hogenhout, TC Mathers, N Pauloski, TA Cernak, JL Wegrzyn, KC Fetter

**Affiliations:** Department of Ecology and Evolutionary Biology, University of Connecticut, Storrs, CT, USA 06269; Department of Medicinal Chemistry, University of Michigan, Ann Arbor, MI, USA 48109; Janelia Research Campus, Howard Hughes Medical Institute, Ashburn, VA, USA, 20147; Department of Crop Genetics, John Innes Centre, Norwich Research Park, Norwich NR4 7UH, UK; Wellcome Sanger Institute: Cambridge, UK; Canton High School, Canton, MI, USA 48187; Department of Molecular and Cell Biology, University of Connecticut, Storrs, CT, USA 06269; Institute for Systems Genomics, University of Connecticut, Storrs, CT, USA 06269; Southeastern Fruit & Tree Nut Research Station, Agriculture Research Service, USDA, Byron GA, USA 31008

**Keywords:** adelgids, reference genome, forest tree health, juvenile hormone, aminopeptidases, invasive species, conservation

## Abstract

Two invasive hemipteran adelgids are associated with widespread damage to several North American conifer species. *Adelges tsugae,* hemlock woolly adelgid, was introduced from Japan and reproduces parthenogenetically in North America, where it has rapidly decimated *Tsuga canadensis* and *Tsuga caroliniana* (the eastern and Carolina hemlocks, respectively). *Adelges abietis*, eastern spruce gall adelgid, introduced from Europe, forms distinctive pineapple-shaped galls on several native spruce species. While not considered a major forest pest, it weakens trees and increases susceptibility to additional stressors. Broad-spectrum insecticides that are often used to control adelgid populations can have off-target impacts on beneficial insects. Whole genome sequencing was performed on both species to aid in development of targeted solutions that may minimize ecological impact. *Adelges abietis* was sequenced using Illumina Linked-Read technology from 30 pooled individuals, with Hi-C scaffolding performed using data from a single individual collected from the same host plant. *Adelges tsugae* used Oxford Nanopore long-read sequencing from pooled nymphs. The assembled *A. tsugae* and *A. abietis* genomes, pooled from several parthenogenetic females, are 220.75 Mbp and 253.16 Mbp, respectively. Each consists of eight autosomal chromosomes, as well as two sex chromosomes (X1/X2), supporting the XX-XO sex determination system. The genomes are over 96% complete based on BUSCO assessment. Genome annotation identified 11,424 and 12,060 protein-coding genes in *A. tsugae and A. abietis*, respectively. Comparative analysis of proteins across 29 hemipteran species and 14 arthropod outgroups identified 31,666 putative gene families. Gene family evolution analysis with CAFE revealed lineage-specific expansions in immune-related aminopeptidases (*ERAP1*) and juvenile hormone binding proteins (*JHBP*), contractions in juvenile hormone acid methyltransferases (*JHAMT*), and conservation of nicotinic acetylcholine receptors (*nAChR*). These genes were explored as candidate families towards a long-term objective of developing adelgid-selective insecticides. Structural comparisons of proteins across seven focal species (*Adelges tsugae*, *Adelges abietis*, *Adelges cooleyi*, *Rhopalosiphum maidis*, *Apis mellifera*, *Danaus plexippus*, and *Drosophila melanogaster*) revealed high conservation of nAChR and ERAP1, while JHAMT exhibited species-specific structural divergence. The potential of JHAMT as a lineage-specific target for pest control was explored through virtual drug and pesticide screening.

## Introduction

Adelgids (*Adelges* and *Pineus* sp.) include at least 65 species of host-specific, sap-feeding hemipterans within Aphidomorpha (Favret et al. 2015). These insects typically exhibit a life strategy similar to their aphid relatives, primarily feeding on plant sap from phloem tissue, yet they are uniquely associated with conifers (Sano and Ozaki 2012). A key distinction to their aphid relatives is the longer, more flexible stylet of adelgids which permits feeding on both vascular and nonvascular tissue (Dancewicz et al. 2021). Additionally, adelgids do not exhibit viviparity, a reproductive trait common in aphids (Havill and Foottit 2007; Chakrabarti 2018). The introduction of non-native adelgid species into North America has resulted in conservation issues for some conifer species. Notable examples include the eastern spruce gall adelgid (*Adelges piceae*) and hemlock woolly adelgid (*Adelges tsugae*) which have devastated fir and hemlock populations, respectively (Ellison et al. 2018; Fei et al. 2019; Zilahi-Balogh et al. 2017).

The eastern spruce gall adelgid (*A. abietis*; Figure 1A) is anholocyclic on spruce (*Picea*) species, consisting of two parthenogenetic generations per year, with fundatrices (asexual females) overwintering at the base of buds. A large number of galls can occur on a single tree causing damage to its crown (Carter 1971) (Figure 1B). Dispersal of *A. abietis* occurs through movement of both winged gallicolae and first instar-nymphs (i.e, crawlers). In North America, *A. abietis* has been observed on black spruce (*P. nigra*), white spruce (*P. glauca*), red spruce (*P. rubens*), as well as non-native European spruces, like the Norway spruce (*P. abies*) (Cook et al. 2012) (Figure 1C). Considered native to Europe and non-native to North America (Plumb 1952), the evolutionary and invasion history of this species remains unclear.

**Figure 1.**
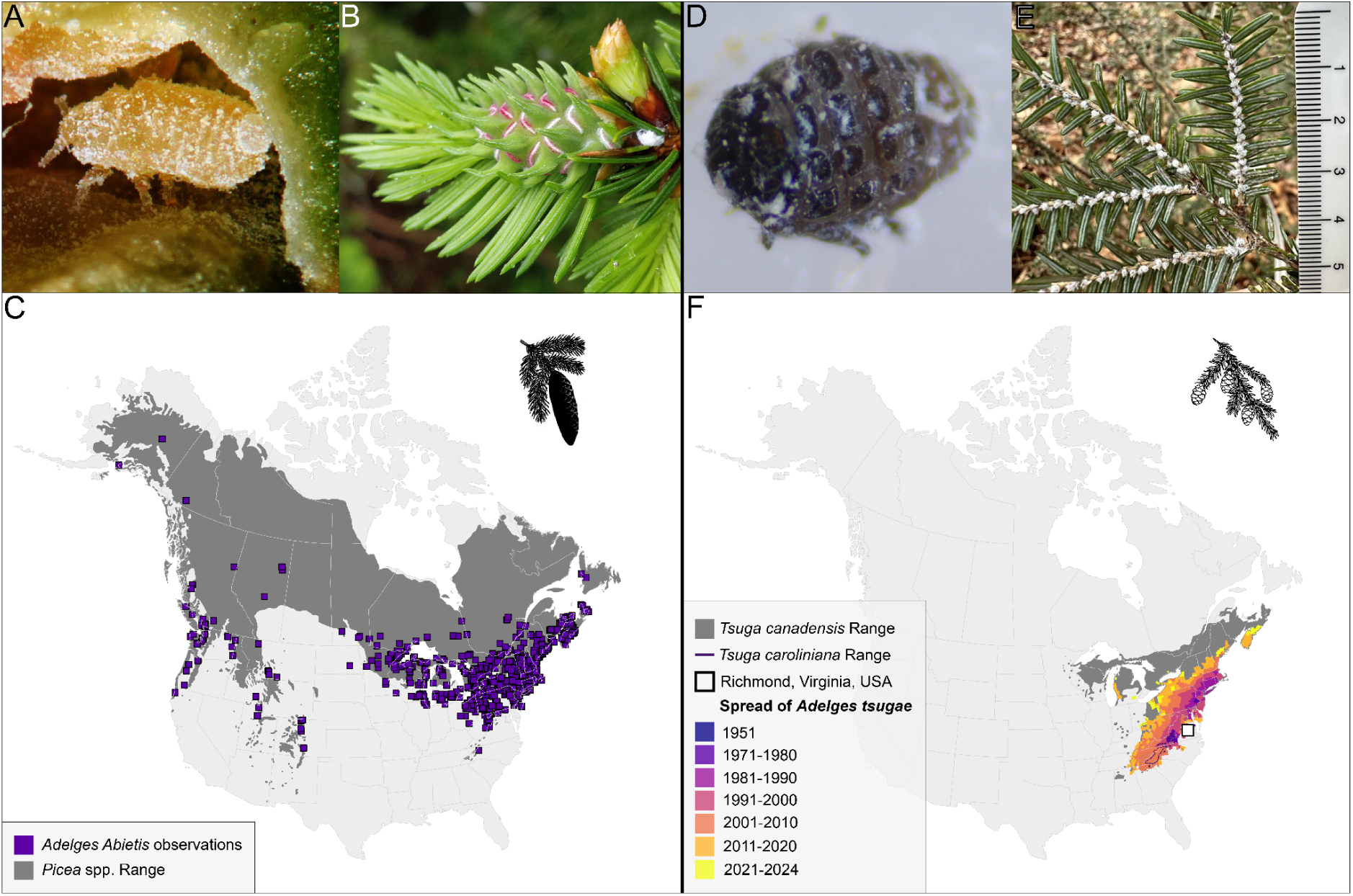
(A) Immature gallicolae of *Adelges abietis* within a gall on *Picea abies* (source: influentialpoints.com). (B) *A. abietis* gall in early development, with stem mother visible. (C) Distribution of North American *Picea* species and observations of *Adelges abietis* (source: GBIF, iNaturalist) - Inset: silhouette representing a spruce. (D) Hemlock woolly adelgid adult. (E) Eastern hemlock branch infested with *A. tsugae* (scale bar in cm). (F) Range of *Tsuga canadensis* and *Tsuga caroliniana* (E. Little and Luther 1971) and the dispersal of *Adelges tsugae* since 1951 (source: 1951-2020 Data derived from USDA FS-FHP, 2021-2024 observations compiled from infestation reports) - Inset: silhouette representing a hemlock.

Evolving in eastern Asia, *A. tsugae* was introduced from Japan into eastern North America during the early 20th century and first collected in Virginia in 1951 (Figure 1D) (Havill et al. 2006a). In its native range, *A. tsugae* alternates hosts between hemlock and spruce (Havil et al. 2016). The invasive *A. tsugae* lineage reproduces parthenogenetically in North America, with endemic spruce species unsuited for sexual reproduction (McClure 1989). HWA disperse as crawlers and are inadvertently moved by humans, wind, birds, and deer (McClure 1990; Russo et al. 2019). While winged morphs are mobile and capable of reproducing in North America, their offspring cannot survive on North American hosts. After settling on the underside of hemlock branches near the base of the petiole, HWA crawlers synthesize white, woolly sacs and remain at their feeding site for the remainder of their life (Montgomery, Bentz, and Olsen 2009) (Figure 1E). The rapid population growth of *A. tsugae* within a single year, combined with its capacity to hitchhike on vertebrates, has enabled it to become a major pest of native eastern North American hemlocks (*Tsuga canadensis* and *Tsuga caroliniana*; Figure 1F), leading to significant ecological and economic threats to forest ecosystems (Orwig and Foster 1998)

Genomic resources in *Adelges* are growing, although the vast majority of adelgids lack a reference genome or other genetic resources (Dial et al. 2023). Mitochondrial sequences from *A. tsugae* and other species determined the invasion history of *A. tsugae*, pinpointing southern Japan as the source of the invasive eastern North American population (Havill et al. 2006b). A study using microsatellites and a mitochondrial marker demonstrated that the HWA populations in western North America are closely related to HWA populations in southern Japan, and likely arrived to western North America through vicariance of the circum-boreal forest during the Plio-Pleiostocene, or possibly via migratory birds (Havill et al. 2016). The fully sequenced *A. tsugae* mitochondrial genome enhanced the phylogenetic resolution of the species and resolved relationships among Aphiodoidea (Havill et al. 2006a; Yeh, Ko, and Wu 2020). Metagenomic sequencing of *A. tsugae* eggs enabled full characterization of the endosymbiont genomes, enabling a better understanding of their contributions to adelgid physiology (Weglarz et al. 2018).

Inspired by the selective targeting of biochemical pathways for human disease with modern precision pharmaceuticals (Chaganti et al. 2024; Juang and Cernak 2025), our hope is that an adelgid-specific pesticide – an adelgicide – could one day be developed as a novel strategy in forest ecosystem management. This study contributes reference genomes for two invasive *Adelges* species with a long-term objective to support the development of targeted pest control strategies. The urgent need for species-specific control methods stems from the ecological damage caused by broad-spectrum insecticides in current use like imidacloprid, which, while effective against adelgids, also harm beneficial insects in part because the nAChR target of these chemicals is conserved across insects. Gene family evolution was investigated across a phylogenetically diverse set of insects to identify lineage-specific expansions and contractions of gene families. This analysis identified gene families involved in juvenile hormone synthesis, neurotransmission, and immune function, which are established or emerging targets for insect pest control. Generative protein structure predictions were employed to illuminate footholds for selectivity in three-dimensional protein-ligand binding pockets, with the goal of identifying features that could be exploited in the development of precision adelgicides.

## Methods & Materials

### DNA Sampling, Extraction, Preparation, and Sequencing

*Adelges abietis* were collected from a single gall growing on *Picea rubens* at Timberline Mountain (39°03’19.6”N 79°23’07.9”W) in Davis, WV, USA on 21 July 2019. Genomic DNA for a 10X Chromium Linked-Read library was extracted from a pooled sample of approximately 30 individuals (detailed methods in File S1). Isolated DNA was quantified on a Qubit 2.0 fluorometer and DNA fragment size was checked by running on a 1.0% agarose gel. The Linked-Read library was prepared following the manufacturer’s protocol and sequenced on an Illumina Nextseq 550. Hi-C Illumina short reads were generated for scaffolding from a single individual collected from a single gall growing on the same *Picea rubens* accession on 16 August 2021. This insect was frozen in liquid nitrogen in an Eppendorf tube, ground with a plastic pestle in 1 mL 1% formaldehyde for 20 minutes. Formaldehyde was quenched by adding 110 μL 1.25 M Glycine for 15 min. Sample was spun at top speed for 15 min in tabletop centrifuge, liquid was removed and replaced with Phosphate Buffered Saline, spun again, the supernatant was removed and sample was stored at –80°C prior to shipping to Phase Genomics (Seattle, WA) for Hi-C library preparation and sequencing.

*Adelges tsugae* were collected from a single accession of *T. canadensis* growing at the Mountain Research Station in Waynesville, NC (35.487 °N, –82.967 °W) in October 2023. Approximately 1.21 m of one branch was collected and stored at 4 °C for 48 hours. Individual adelgids were collected into Macherey-Nagel Lysis Buffer T1 using forceps and a dissection microscope. A total of 400 sistens (adelgid nymphs) were collected to provide sufficient material for DNA extraction. DNA was extracted from the pooled sample using the Qiagen MagAttract high molecular weight (HMW) DNA Kit, with modifications based on (Mao et al. 2023) and further adjustments to timings, temperatures, and agitation intensities (File S1). Isolated DNA quality was measured with Thermo Scientific NanoDrop One spectrophotometry, Thermo Scientific Qubit 4 DNA fluorometry, and Agilent 4200 TapeStation Genomic DNA electrophoresis. Oxford Nanopore Technologies Ligation Sequencing Kit v14 (SQK-LSK114) was used to prepare the library. DNA repair was performed at 20 °C for 20 min followed by 10 min at 65 °C to inactivate the enzymes. DNA was eluted at 37 °C. The final library was quantified using a Qubit 4 fluorometer. The flow cell was loaded three times each with 18 fmol of library.

### RNA Sampling, Extraction, Preparation, and Sequencing

Individual *Adelges tsugae* were collected from branches of *T. canadensis* growing in Leesburg, VA, USA (39°07’05.3”N 77°34’07.5”W) on 6 February 2024, and *A. abietis* were collected from galls in Davis, WV (39°03’19.6”N 79°23’07.9”W) on 22 August 2019 as described above. Salivary glands were dissected by placing insects in phosphate buffered saline and then grabbing insects with forceps at the pronotum and anterior abdomen and pulling apart. The salivary glands remain attached to the brain on the anterior portion. The glands were isolated from other tissue using Minutien insect pins, with their tips bent approximately 30 degrees, mounted in micro dissecting needle holders. Separated glands and the remaining carcass were collected separately into 100 μL of Arcturus PicoPure Extraction buffer. Total RNA was prepared using the Arcturus PicoPure RNA Isolation kit including the optional DNAse step. Barcoded RNASeq libraries were prepared for Illumina NextSeq 550 sequencing (150bp PE) with a method described previously (Cembrowski et al. 2018). Libraries with fewer than 15 million reads or mean read length below 70 bp were excluded as evidence for structural annotation (Table S1).

### Genome assembly

#### Adelges abietis

A total of 1.19 Gb of 10X linked reads were assembled into a draft genome with Supernova with default parameters. Hi-C reads were aligned to the *Supernova* draft assembly with Juicer v1.6.2 (Durand et al. 2016). The 3D-DNA pipeline (Dudchenko et al. 2017) was applied with default settings in “haploid mode” to correct mis-assemblies and generate chromosome-scale super scaffolds. Manual review of the 3D-DNA assembly was conducted with Juicebox Assembly Tools (Durand et al. 2016). The scaffolded assembly was screened for contamination with BlobTools v1.0.1 (Laetsch and Blaxter 2017; Sujai Kumar et al. 2013) using aligned 10X Genomics linked reads and taxonomy information from BLASTN v2.2.31 (Camacho et al. 2009) searches against the National Center for Biotechnology Information (NCBI) nucleotide database (nt, downloaded October 13, 2017). To calculate per-scaffold coverage, 10X Genomics linked reads were debarcoded with process_10xReads.py from the proc10xG package (https://github.com/ucdavis-bioinformatics/proc10xG) and aligned to the reviewed 3D-DNA assembly with BWA mem v0.7.7 (Vasimuddin et al. 2019). Finally, the decontaminated assembly was sorted by chromosome length, and assessed for completeness with BUSCO v5.4.5 (Manni et al. 2021), QUAST v5.2.0 (Gurevich et al. 2013), and by comparing K-mer content of the debarcoded 10X Genomics linked reads to the assembly with KAT comp v2.3.1(Mapleson et al. 2017; Shen, Sipos, and Zhao 2024). The genome was assessed for accuracy with Merqury v1.3 (Rhie et al. 2020).

#### Adelges tsugae

Nanopore reads were basecalled from a single PromethION R10.4.1 flow cell running MinKnow v23.07.12. Initial statistics on the read count, length, and base quality were assessed using NanoPlot v1.33.0 (De Coster and Rademakers 2023). Reads passing an initial quality cutoff of >Q10 were screened for DNA contamination against RefSeq genomes (release 221) of archaea, bacteria, fungi, plants, and viruses using Centrifuge v1.0.4-beta and reassessed using NanoPlot after removal of contaminant-classified reads (D. Kim et al. 2016). The estimated genome size was obtained via kmer count using kmerfreq v4.0 (Hengchao Wang et al. 2020) with GCE v1.0.2 (Hengchao Wang et al. 2020). Filtered reads were assembled using both Flye v2.9.1 (Kolmogorov et al. 2019) and Canu v2.2 (Koren et al. 2017). Initial draft assemblies and each subsequent iteration were assessed using BUSCO hemiptera_odb10 for completeness, Merqury v1.3 (Rhie et al. 2020) for quality, and QUAST for contiguity. The Canu assembly was selected for downstream analysis. Heterozygosity was reduced for draft assemblies using Purge Haplotigs v1.1.2 (Roach, Schmidt, and Borneman 2018). The contig-level assemblies were scaffolded against the chromosome-level *A. abietis* genome using RagTag v2.1.0 (Alonge et al. 2022), reordering contigs into putative chromosomes, in descending order by length. Following scaffolding, annotation transfer between genome assemblies was performed using LiftOff v1.6.3 (Shumate and Salzberg 2022).The final assembly was re-assessed for quality metrics.

The *Adelges tsugae* and *A. abietis* final assemblies were screened for contaminant and adaptor sequences using NCBI Foreign Contamination Screen (FCS-GX and FCS-adaptor) v0.5.4 (Astashyn et al. 2024).

### Structural and functional annotation

The repeat library for each species was generated *de novo* via RepeatModeler v2.0.4 (Flynn et al. 2020). This library was used to softmask the assembled references with RepeatMasker v4.1.4 (Smit, AFA, Hubley, R & Green, P. 2013-2015). The softmasked reference genomes were provided to EASEL (https://gitlab.com/PlantGenomicsLab/easel) v1.5 (regressor set to 70) to facilitate the structural and functional identification of protein-coding genes (Hart et al. 2020). Transcriptomic evidence provided gene prediction support within EASEL through the alignment of RNASeq reads to their corresponding genome. The *Adelges tsugae* annotation was informed by four salivary gland libraries, four carcass libraries (salivary glands removed) (SRR30936309), and an adult overwintering sistens library (SRR1198669). The *Adelges abietis* annotation was informed by two salivary gland libraries, four carcass libraries, and a whole body library (SRR30936310). Additional protein model support for annotation was provided by NCBI’s RefSeq protein database (release 208) (Goldfarb et al. 2025) and OrthoDB v11 (Kuznetsov et al. 2023).

Genome synteny was assessed using protein-protein comparisons of the *Adelges tsugae*, *A. abietis*, *A. cooleyi* (using the top 10 scaffolds, following the approach of (Li et al. 2023), and *Daktulosphaira vitifoliae* annotations using MCScanX (Wang et al. 2012), and was visual with SynVisio (Bandi and Gutwin 2020).

### Comparative genomics among hemipteran

All hemipteran species with genome annotations were downloaded on 17 October 2024 from NCBI (45 total) and were considered for comparative analysis, and 14 related species were used as outgroups (Table S2) (Bailey et al. 2022; Panfilio et al. 2019; Ma et al. 2021). All protein sets had a BUSCO completeness above 80% and no obvious anomalies in regards to total gene count. Orthofinder v2.5.5 was run with 45 species to classify potential orthologous genes and orthogroups with default settings: MAFFT for alignments, FastTree for maximum likelihood gene trees, and the STAG consensus method for species tree inference (Emms and Kelly 2019). Clade support in the species tree reflects the proportion of orthogroup gene trees supporting each branch. The longest sequence from each orthogroup was used as a representative to assign a function with EnTAP v1.2.1 (Hart et al. 2020) that integrated EggNOG v5.0.2 (Huerta-Cepas et al. 2019), Refseq Invertebrate release 224, and Swiss-Prot (release 2024_03).

Significant orthogroup expansions were identified with Cafe v5.1 (Mendes et al. 2021) based on birth and death process models. Gene turnover was estimated using the maximum likelihood inference method. The distances in the rooted tree obtained from the single-copy genes (from OrthoFinder) were transformed into ultrametric units (Table S3). Orthogroups with representation in only one species (CAFE5 clade_filter), overrepresentation in one species (one species ≥50 orthologs and all others ≤3), and extreme differentials between minimum and maximum counts (63 or more orthologs), were removed. Gene family expansion or contraction was considered significant at *P* < 0.01.

### Assessing significantly expanded/contracted gene families

Hidden Markov models (HMMs) for Juvenile hormone (JH) acid methyltransferase (*JHAMT*; PTHR43464), Survival of motor neuron (SMN; PTHR39267), Endoplasmic reticulum aminopeptidase protein 1 (*ERAP1*; PTHR11533), Juvenile hormone binding protein (*JHBP*; PF06585), and nicotinic acetylcholine receptor (nAChR) alpha-7 subunit were retrieved from PantherDB (v17.0; (Thomas et al. 2022)) and Pfam (v35.0; (Finn et al. 2014)), cross-referenced through InterProScan (v5.59-91.0; (Jones et al. 2014)) to confirm domain architectures. HMM profiles were processed into searchable databases using hmmpress (HMMER v3.3.2; (Z. Zhang and Wood 2003)) with default parameters. Subsequent batch CD-Search analysis (NCBI CDD v3.20; (Marchler-Bauer et al. 2015)) validated domain presence across targets using RPS-BLAST (E-value <1e−10, coverage >70%). Validated sequences were aligned with MAFFT v7.526 (L-INS-i algorithm; BLOSUM62 matrix, gap penalty=1.53; (Katoh and Toh 2008)) via the Computational Biology Research Consortium (CBRC) web server, followed by refinement with MaxAlign v1.3 (Gouveia-Oliveira et al. 2007) to retain sequences covering ≥75% of consensus length and columns with ≤40% gaps. Genes will be referred to in italics (i.e, *ERAP1, JHAMT*), and protein names by the non-italicized version (i.e, ERAP1, JHAMT).

### Homology modeling and structural comparison

Structures of insect proteins were predicted using AlphaFold 3 (AF3) (Abramson et al. 2024). ERAP and JHAMT proteins were each folded as monomers. As nAChR proteins are pentameric, their structure prediction was simplified by folding all nAChR proteins as homopentamers, although only some nAChR subtypes are known to homopentamerize in vivo (Barrantes 2023). Models were evaluated for quality using their accompanying predicted template modeling (pTM) confidence scores. Proteins whose resulting models had a pTM score below 0.75 were excluded from the analysis in order to prevent the data from being skewed by outlier sequences of abnormally low homology. TM scores were calculated using Zhang et al’s TM align algorithm and the scores normalized to the average length of the pair of compared proteins (Y. Zhang and Skolnick 2004; Xu and Zhang 2010). Scores were then sorted by in-group (adelgids; *A. tsugae*, *A*. *abietis*, *A. cooleyi*) and outgroup (other insects; *R. maidis*, *A. mellifera*, *D. plexippus*, *D. melanogaster*) and arrayed combinatorially for visualization via heatmap.

### Sequence similarity analysis

Protein sequences (i.e. amino acids) were aligned in MEGA v11.0.13 (Sudhir Kumar et al. 2018) with MUSCLE v3.8.31 (Edgar 2004). Amino acid sequence similarities were calculated with the “Sequence Manipulation Suite” for JavaScript using the “Ident and Sim” tool, with similar amino acid groups set to GAVLI, FYW, CM, ST, KRH, DENQ, P (Stothard 2000). Proteins resulting in models with low pTM confidence scores were excluded as mentioned above. Percent similarity scores were then sorted by in-group (adelgids; *A. tsugae*, *A*. *abietis*, *A. cooleyi*) and outgroup (other insects; *R. maidis*, *A. mellifera*, *D. plexippus*, *D. melanogaster*) and visualized via a kernel density estimate (KDE) plot (Table S4).

### Binding site analysis

To identify insect protein binding site residues, full protein sequences for all selected insects were compared to the sequences of homologous proteins from other species with crystal structure data. nAChR sequences were aligned and searched for the conserved active site loop regions, which have been previously reported (Gharpure, Noviello, and Hibbs 2020). JHAMT sequences were aligned to those from three species of moth (*B*. *mori*, *M*. *sexta*, and *O*. *furnacalis*) and identified residues matched to the substrate contacting residues and cofactor contacting residues from the *B*. *mori* crystal structure [Protein Data Bank (PDB) ID: 7EC0, (Guo et al. 2021)]. ERAP1 sequences were aligned with a human ERAP1 sequence with a known crystal structure [PDB ID: 6M8P, (Maben et al. 2021)]. Active site residues were identified by alignment in the crystal structure. C-terminal selectivity region residues were identified by alignment with residues implicated in substrate selectivity (Giastas et al. 2019). The resulting sequences were then compiled and visualized as logo plots (Crooks et al. 2004). Full sequences for ERAP1, JHAMT, and nAChR can be found in Table S4. Isolated binding site sequences for ERAP1, JHAMT, and nAChR can be found in Table S5, Table S6, and Table S7, respectively. Isolated active site sequences for ERAP, JHAMT, and nAChR can be found in Table S8.

### Molecular docking

Homology models generated using AlphaFold3 were prepared for molecular docking using mgltools (version 1.5.7), while ligands were generated by rdkit (version 2024.03.1) and mgltools [(Morris et al. 2009); RDKit: Open-source cheminformatics. https://www.rdkit.org]. The molecular docking was performed on the homology model using the SMINA docking program with the binding pocket centered on coordinates [2.5, –26.5, 41.32] and a pocket size of 20Å × 20Å × 20Å (Koes et al. 2013). The molecular libraries for docking were obtained from PubChem and DrugBank (Knox et al. 2024; Kim et al. 2025). KDE analysis was performed using the seaborn python library (version 0.13.2) and the docking score of –4 kcal/mol was used as a cutoff (Waskom 2021).

## Results and Discussion

### Genome sequencing and assembly

*Adelges abietis* 10X sequencing generated 119M Illumina short reads (Table S9). The Supernova assembled genome produced a reference of 290.32 Mb in 9,607 contigs with an N50 of 4.77 Mb. Hi-C scaffolding of the *A. abietis* genome initially produced an assembly 253.17 Mb in length with 2998.36 N’s per 100 Kb, an N50 of 27.91 Mb, and an accuracy value (QV) of 50.87 (Table S10). However, manual curation revealed structural inconsistencies within Chromosome 1 (signature of translocations in the two haplotypes). Manual adjustments, supported by HiC reads, resolved the assembly into ten chromosomes (Figure S1). Post manual corrections, the assembly was 253.17 Mb in length, with 2884.06 N’s per 100 Kb, an N50 of 21.97 Mb, and the QV remained at 50.87. A total of 87.17% of sequences were contained within the ten chromosomes (eight autosomal and two sex chromosomes). BUSCO assessment found 2471 of 2510 expected complete (C:97.2%) single-copy orthologs from the hemiptera_odb10 database and 47 duplicated (D:1.3%), leaving only 8 fragmented (F:0.3%) and 31 missing (M:1.2%) (Table 1). Contaminant screening of *Adelges abietis* post assembly removed 61 (2,740,304 bp) unplaced scaffolds, of which 54 were classified as *Candidatus Vallotia tarda* (33), *Ca. Vallotia lariciata* (20), *Ca. Profftia tarda* (1), or *Ca*. *Arsenophonus triatominarum* (0.04%) (Table S11). These bacteria represent known endosymbionts that commonly appear as contaminants in adelgid genome assemblies. *Ca. Vallotia tarda* and *Ca. Vallotia lariciata* are betaproteobacterual obligate symbionts found in adelgids, with *Ca. Vallotia tarda* specifically associated with *A. larcis/A. tardus* species complex, and *Ca. Vallotia lariciata* occurring in *A. lariciatus* (Szabó et al. 2022). *Ca. Profftia tarda* is a gammaproteobacterial symbiont that complements *Ca. Vallotia* in essential amino acid biosynthesis and is restricted to larch-associated adelgid lineages (Szabó et al. 2022). *Ca. Arsenophonus triatominarum* is a gamma-proteobacterium originally identified from triatomine bugs (*Triatoma infestans*) and represents a member of the *Arsenophonus* clade known for diverse symbiotic interactions with arthropods (Hypsa and Dale 1997; Martin Říhová et al. 2023).

**Table 1.** Assembly and Annotation Statistics for *Adelges tsugae* and *Adelges abietis*.

| **Assembly** | **Total Chromosomes** | **N50 (Mb)** | **Length (bp)** | **Sequence in Chromosome** | **% Gaps** | **% Repeat** |
| --- | --- | --- | --- | --- | --- | --- |
| *Adelges tsugae* | 8+2 | 21.14 Mb | 221,267,597 | 95.49% | 2.402% | 17.90% |
| *Adelges abietis* | 8+2 | 21.97 Mb | 253,164,600 | 87.2% | 2.884% | 24.91% |
| **BUSCO Genome, hemiptera** |  |  |  |  |  |  |
|  | **Complete** | **Single** | **Duplicate** | **Fragment** | **Missing** | ***n* Searched** |
| *Adelges tsugae* | 99.3% | 98.2% | 1.1% | 0.4% | 0.3% | 2510 |
| *Adelges abietis* | 98.5% | 97.2% | 1.3% | 0.3% | 1.2% | 2510 |
| **BUSCO Annotation, hemiptera** |  |  |  |  |  |  |
|  | **Complete** | **Single** | **Duplicate** | **Fragment** | **Missing** | ***n* Searched** |
| *Adelges tsugae* | 98.0% | 96.7% | 1.3% | 0.2% | 1.8% | 2510 |
| *Adelges abietis* | 95.9% | 94.7% | 1.2% | 0.2% | 3.9% | 2510 |

*Adelges tsugae* was collected from several branches of trees in Leesburg, VA. Although *A. tsugae* samples for DNA extraction were collected elsewhere in Waynesville, NC, it is unlikely that there is a high level of genetic differentiation between individuals from these locations, since in eastern North America, *A. tsugae* reproduces only parthenogenetically. The clonal diversity of this population as determined by the ratio of the number of determined clonal multilocus lineages to the number of assessed individuals was determined to be very low, with a value of 0.033 (Havill et al. 2016). *A. tsugae* sequencing generated 100.78 Gb of long reads (ONT) with an 11.02 Kb N50, of which 89.94 Gb passed an initial Q10 quality score threshold and 77.71 Gb passed contaminant screening (File S2; Table S9). Among the 4,037,933 (13.60%) reads classified as contaminants, the most abundant species was *Candidatus Pseudomonas adelgestsugas*, one of *A. tsugae*’s two dual-obligate endosymbiotic bacteria, with just over 3M unique reads representing 75.7% of those removed. The other paired endosymbiont, *Ca. Annandia adelgestsuga*, was not specifically identified. However, sequences with closest taxonomic matches to *Ca. Annandia pinicola* (4.43%) and 19 intraspecific variants of *Buchnera aphidicola* (together 0.59%) were identified. Given the similarity of the *Buchnera* and *Annandia* genomes, as well as the similarity-based metagenomic classification technique used, these reads may represent *Ca. Annandia adelgestsuga* (Weglarz et al. 2018). The second most abundant classification was *Serratia symbiotica* (9.05%), an endosymbiont unique to eastern North America and *Tsuga sieboldii*-specific *A. tsugae* lineages in Japan (C. D. von Dohlen et al. 2013). This was followed by *Porphyrobacter* sp. GA68 (0.66%), *Cellulophaga* sp. HaHaR_3_176 (0.18%), and *Rhizoctonia solani* (0.14%), an indoor air bacterium, a marine bacterium, and a plant-pathogenic fungus, respectively (Table S12).

The genome size of *Adelges tsugae* was estimated at 267.97 Mb yielding 290x read coverage. The Canu assembled draft genome contained 2,131 contigs with a 7 Mb N50. Reducing heterozygosity yielded 377 contigs with an N50 of 9 Mb. Scaffolding against the chromosome scale *A. abietis* reference yielded 345 contigs with a 21 Mb N50. This final assembly is 221.26 Mb in length, 95.49% of which is contained within ten putative chromosomes (eight autosomes and two sex chromosomes). This reference reports 1.45 N’s per 100 kb, an N50 of 21.14 Mb, reflecting exceptional continuity. The Merqury QV score of 42.88 indicates a base-level accuracy of 99.99%, further underscoring the high quality of the assembly. BUSCO assessment found 2464 of 2510 expected complete (C:99.3%) single-copy orthologs from the hemiptera_odb10 database and 28 duplicated (D:1.1%), leaving only 10 fragmented (F:0.4%) and 8 missing (M:0.3%) (Table 1).

Of the assembled Aphidomorpha genomes, Adelgidae and Phylloxeridae tend to have smaller genomes than the Aphididae. (Table S2) While Aphididae genomes are variable in size, ranging between 300-600 Mb (Huang et al. 2025), the average genome size of the characterized Aphididae is 369.4 Mb. In contrast, all of the assessed Adelgidae and Phylloxeridae genomes are smaller than 300 Mb. *Adelges tsugae* is the smallest, with a genome size of 220.75 Mb, and *A. abietis*, *A. cooleyi*, and *Daktulosphaira vitifoliae* have estimated genome sizes of 253.16 Mb, 270.2 Mb, and 282.6 Mb, respectively (Rispe et al. 2020; Dial et al. 2023).

### RNA sequencing and quality control

For *Adelges tsugae*, eight RNASeq libraries passed quality control (150bp PE), ranging from 18.1 to 25.3M reads per library. For *A. abietis*, seven RNASeq libraries passed quality control (150bp PE), ranging from 15.5 to 24.4M reads per library (Table S1).

### Structural and functional annotation

A total of 17.66% of the *Adelges tsugae* genome and 28.31% in *A. abietis* were softmasked and identified as repetitive DNA (Table 1; Table S13; Figure 2A, 2B). The overall repeat content masked in both *A. abietis* and *A. tsugae* are lower than the repeat content of *A. cooleyi* (43.6% of genome masked; (Dial et al. 2023)) and the phylloxeran *Daktulosphaira vitifoliae* (53.26% of the genome masked; (Z. Li et al. 2023)), on the other hand, the range of repeats masked in the adelgid genomes presented here fall within that of aphids, *Myzus persicae* (22% of genome masked), and *Acyrthosiphon pisum* (29% of genome masked; (Mathers et al. 2021)). LINEs are the most abundant elements in both genomes (2.22% *A. abietis*, 2.75% *A. tsugae*) while LTRs (1.04% *A. abietis*, 0.23% *A. tsugae*) and SINEs (0.00% *A. abietis*, 0.01% *A. tsugae*) are less common, a pattern common among some hemipterans including aphids and whiteflies (Petersen et al. 2019). *Adelges tsugae* differs from the pattern in DNA transposon content with only 0.46% of sequence compared to *A. abietis*’s 2.14%, though this and the overall low TE abundance may be explained by the large portion of unclassified TEs (*A. abietis* 20.59%, *A. tsugae* 11.68%). Protein-coding genes show variable densities across the genome and do not support a consistent positional pattern, with both gene-rich and gene-poor regions distributed throughout the chromosome arms rather than concentrated at specific locations (Figure 2C, 2D) The repeat age distribution of *A. tsugae* and *A. abietis* reveals a continuous pattern, marked by distinct proliferation phases (Figure 2A, 2B). The most recent proliferation phase in *A. tsugae* is characterized predominantly by DNA transposons and LINE elements (Kimura distance <4; Fig 2A and Fig 2B), with minor contributions from LTR retrotransposons. Earlier activity shows comparable profiles dominated by unclassified elements, though LINEs and DNA transposons maintain consistent representation across all phases (Kimura distance > 4, < 7). Notably, while LTR retrotransposons constitute a smaller fraction of each burst, they persist through all proliferation phases. The TE profile in *A. tsugae* differs from *A. abietis* and other aphids like *Myzus persicae* and *Metopolophium dirhodum*, which display recent expansions dominated by DNA transposons (Figure 2A, 2B; Table S13) (Baril et al. 2023).

**Figure 2.**
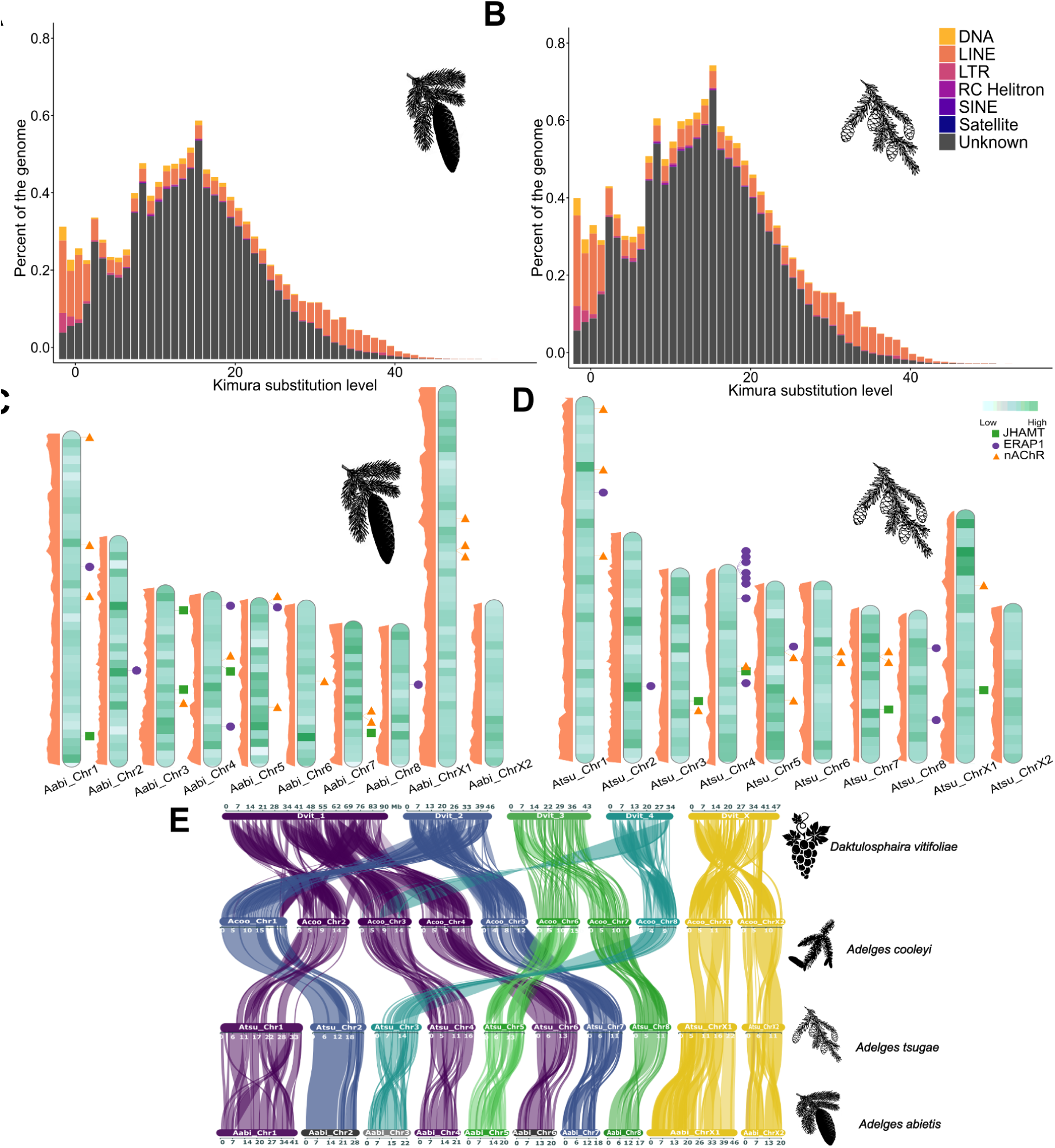
(A,B) Repeat landscapes of known transposable element (TE) families (DNA transposons, LINEs, and LTR retrotransposons) and unclassified repeat families in *Adelges tsugae* and *A. abietis*, respectively. (C,D) Chromosome models of *A. tsugae* (left) and *A. abietis* (right) with gene density from light to dark shown in green (*A. tsugae* 0-56 genes per 500,000 bp, *A. abietis* 1-49). The focal gene family members, including: *JHAMT* (green boxes)*, ERAP* (purple circle), and *nAChR* (orange triangle) are shown in their respective positions on the far right. The repeat density track is on the left side in orange. (E) Synteny analysis between *Daktulosphaira vitifoliae*, *Adelges cooleyi* (longest ten scaffolds), and the ten pseudo-chromosomes of *A. abietis* and *A. tsugae*. Adelgids exhibit strong synteny in both sex chromosomes and autosomes.

Identification of protein-coding regions in *Adelges tsugae* resulted in 11,424 protein-coding genes with BUSCO completeness of C:98.0% [S:96.7%, D:1.3%], F:0.2%, M:1.8%. A total of 12,060 *A. abietis* protein-coding genes were identified with a BUSCO completeness of C:95.9% [S:94.7%, D:1.2%], F:0.2%, M:3.9% (Table 1;Table S14; Table S15). Functional annotations, including both sequence similarity searches and alignments to EggNOG gene families, were available for 94.32% of *A. tsugae* protein-coding genes and 90.69% of *A. abietis* genes (Table S15).

Synteny analysis revealed strong conservation of chromosomal architecture across adelgid genomes, with *A. cooleyi*, *A. tsugae* and *A. abietis* supporting a structure of eight autosomes and two sex chromosomes (X1 and X2) (Figure 2E, Table S16). Adelgids, phylloxera, and aphids largely share an XX-XO sex chromosome system. Despite this, their chromosomes exhibit different evolutionary patterns. Recent comparative studies by Li et al (Z. Li et al. 2023) observed frequent inter-chromosomal autosomal rearrangements (and frequent intra-X-chromosomal rearrangements) among *D. vitifoliae*, *A. cooleyi*, and *A. pisum*, and suggested that the elevated rate of autosomal rearrangements is a lineage-specific feature of aphids. However, Huang et al (Huang et al. 2025), with a broader sampling, noted more variation among aphid lineages, and specifically that those less speciose had fewer inter-autosomal rearrangements. Among the three genomes representing adelgids, there appears to be fewer autosomal and X chromosome rearrangements. In aphids, Li et al (Y. Li, Zhang, and Moran 2020) observed that the X chromosomes were characterized by TE expansions, and genes with low levels of expression and under weak purifying selection. Among the assembled adelgids, X1 of *A. abietis* was both larger and more repeat rich (45.9 Mb, 2,505 genes, 21% of the gene space) compared to *A. tsugae* (26.9 Mb, 1,504 genes, 13% of the gene space) (Table S17).

### Comparative genomics among hemipteran

Comparative genomic analysis across hemiptera identified 32,340 orthogroups comprising 93.69% of total proteins examined, of which 85.61% had functional annotations (Table S3). Of the 1,908 orthogroups shared across all 45 species, only 59 were single-copy orthologs. While most orthogroups (68.93%) lacked adelgid representation, 862 were adelgid-specific, with 205 shared across all three *Adelges* species. A total of 33, 189, and 123 groups were unique to *A. tsugae*, *A. abietis*, and *A. cooleyi*, respectively (Figure 3A).

**Figure 3.**
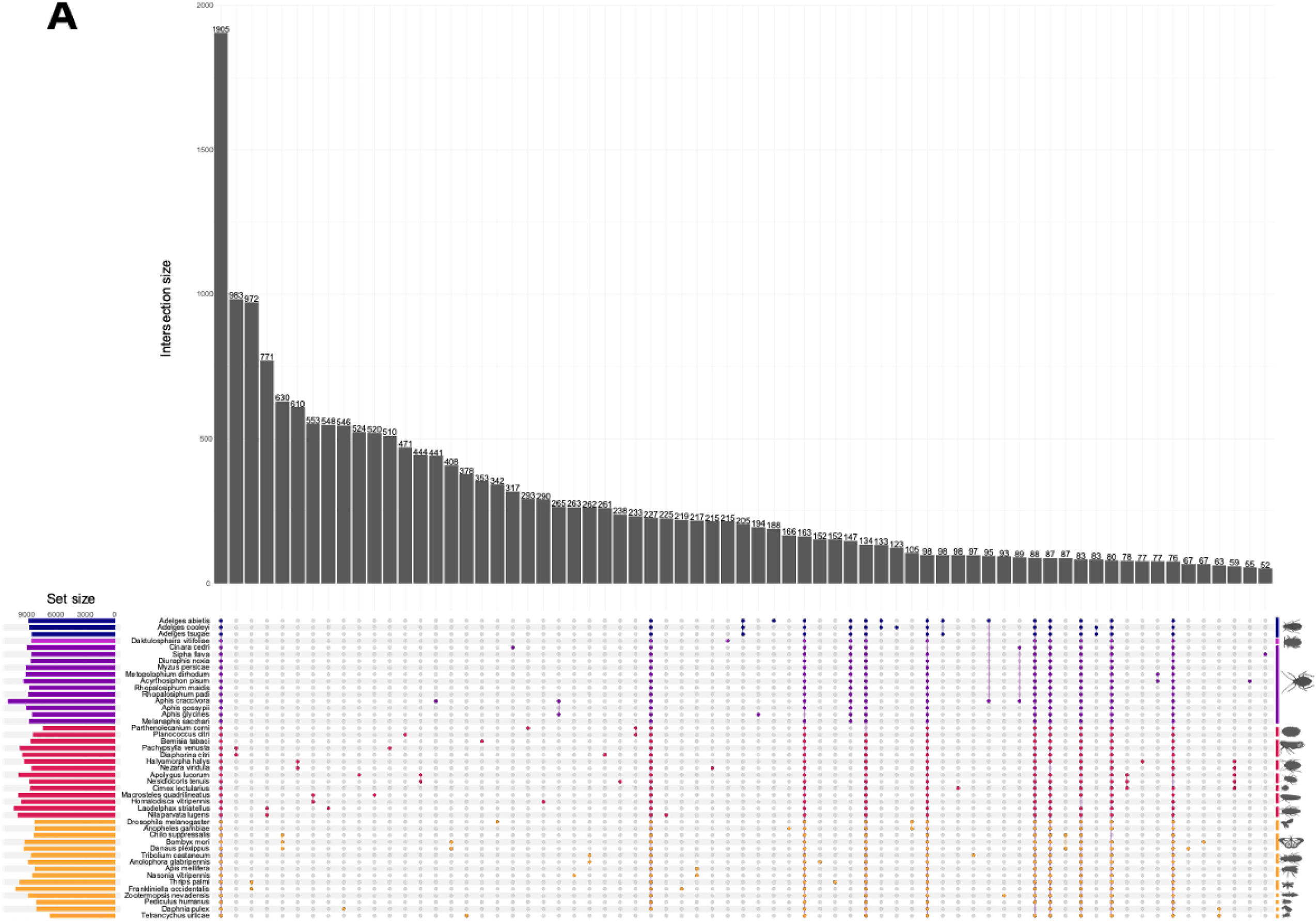
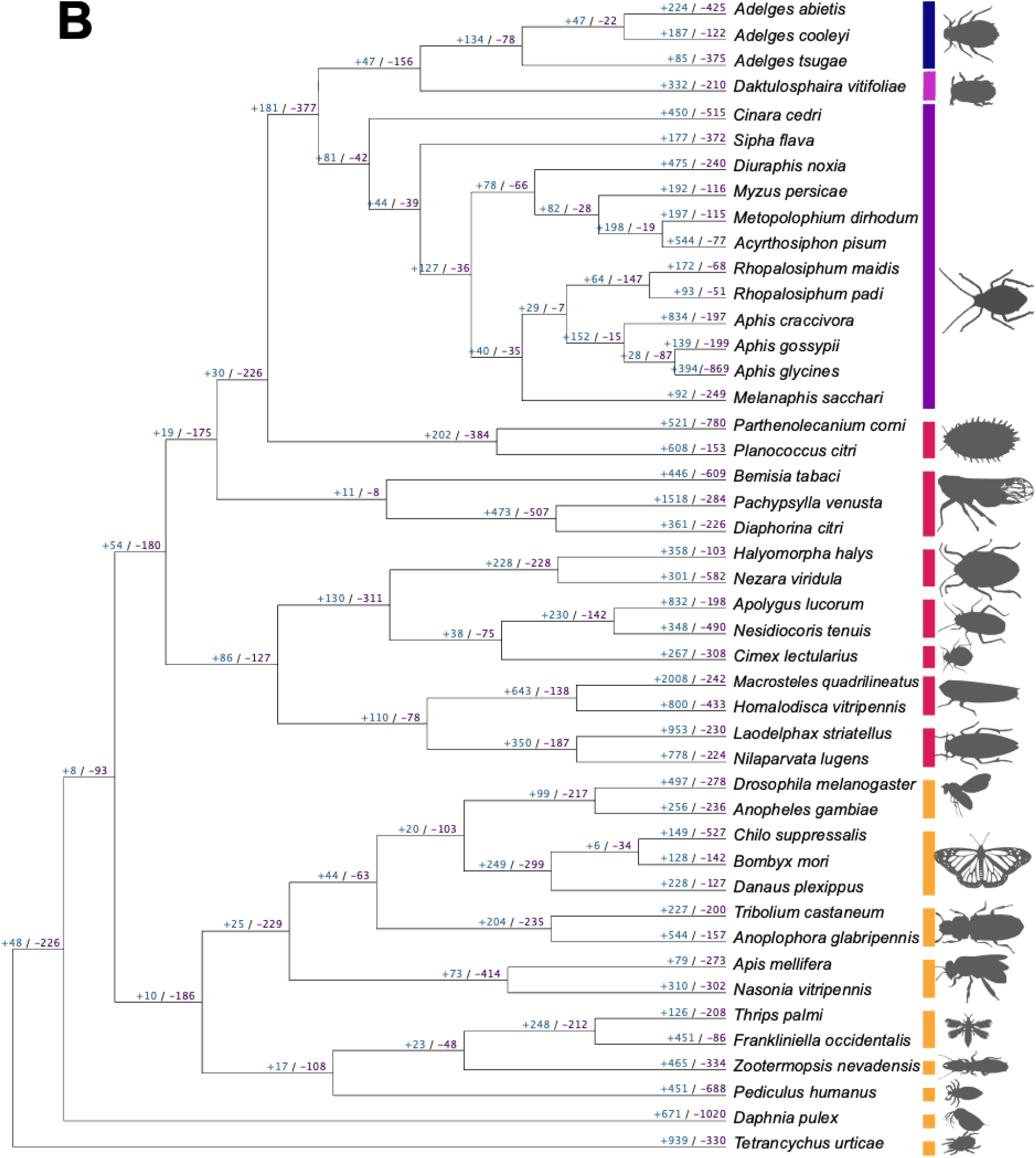
(A) Orthologous gene group interactions between 45 species. Minimum interaction size displayed is 50. Rightmost vertical bars represent lineages, and are grouped from top to bottom and darkest to lightest: Adelgidae, Phylloxeridae, Aphididae, other hemiptera, and non-hemipteran arthropods. (B) Phylogenetic tree (ultrametric) was generated with Orthofiner with the number of significantly expanding and contracting genes indicated at nodes. Silhouette images from bugwood.org.

Phylogenetic analysis of 45 high-quality genomes found 244 gene families with significant changes in size (Table S18; Figure 3B). Comparison of gene family dynamics identified 235 significantly divergent orthogroups after filtering transposable elements (Table S19). Of these, 32 orthogroups were absent across all *Adelges*, with 31 likely lost in the adelgid-phylloxeran common ancestor (Table S20). Among lost gene families, signal transduction mechanisms were most abundant, including G-protein receptors, immunoglobulins, and PET domain proteins.

At the *Adelges* most recent common ancestor (MRCA) node, 24 orthogroups are expanding and 34 contracting. Within this lineage, *A. tsugae* has 19 expanding and 62 contracting orthogroups, *A. cooleyi* has 42 expanding and 27 contracting, and *A. abietis* has 29 expanding and 46 contracting. The largest expansions in *A. abietis* were associated with carbohydrate transport and metabolism, particularly ABC transporters and carboxypeptidases (Table S21). This suggests adaptation to host plant metabolism, a pattern commonly observed in specialized herbivores coping with host plant defenses (Y.-T. Wu et al. 2024; C. Wu et al. 2019). While phylloxera evolved genes for host interaction, adelgids expanded core metabolic functions, focusing on energy production (Rispe et al. 2020). This pattern aligns with highly specialized herbivores, showcasing distinct adaptive strategies between closely related species (C. D. V. O. N. Dohlen and Moran 2000).

While the comparative analysis revealed broad patterns of gene family evolution in Adelgidae, a closer examination of specific gene families provides insights into potentially adaptive mechanisms critical for host interaction and survival. Genes involved in the suppression of host immune recognition, regulation of developmental processes, and signal transduction pathways are of particular interest because they likely contribute to the insect’s ability to evade host defenses, adapt developmentally to environmental cues, and coordinate physiological responses essential for successful parasitism and reproduction. The following families expanded below were selected as they were shown to be rapidly expanding/contracting with respect to the most recent common ancestor of the adelgids, *A. tsugae*, *A. abietis*, and *A. cooleyi*. In addition, nAChR was assessed as a canonical targer that lacked a unique trend for adelgids

### Signal transduction gene families

As members of the cysteine (Cys)-Loop ligand-gated ion channel (CysLIGC) receptor family, *nAChR*s are highly conserved across insects and most animals (Marcovich et al. 2020). nAChRs are composed of five subunits, being either homopentameric or heteropentameric, and are membrane-bound, with extracellular, transmembrane, and intracellular regions (Figure 4A). Insect nAChR subunits can be separated into two distinct classes, α and β, which are primarily differentiated by the presence or absence of a disulfide linkage between two adjacent cysteine (Cys) residues in the ligand binding site. The ligand binding site is located in the extracellular region at the interface between two subunits, and is composed of six loops, with three being provided by each subunit in the interface. In order to bind a ligand, at least one of the subunits at the given interface must be an 𝛼 subunit, with the other being either α or β (Rosenthal and Yuan 2021). Since nAChRs are the target of imidacloprid, the most commonly used insect control methods for adelgids (McCarty and Addesso 2019), a better understanding of *nAChR* diversity within Adelgidae and between Adelgidae and Insecta was sought. While *nAChR* composition and expression is not the only factor determining susceptibility to chemical controls, expression of different subunits and mutations within subunits can impact susceptibility to nicotinic compounds (Zhang et al. 2018; Cartereau et al. 2020).

**Figure 4.**
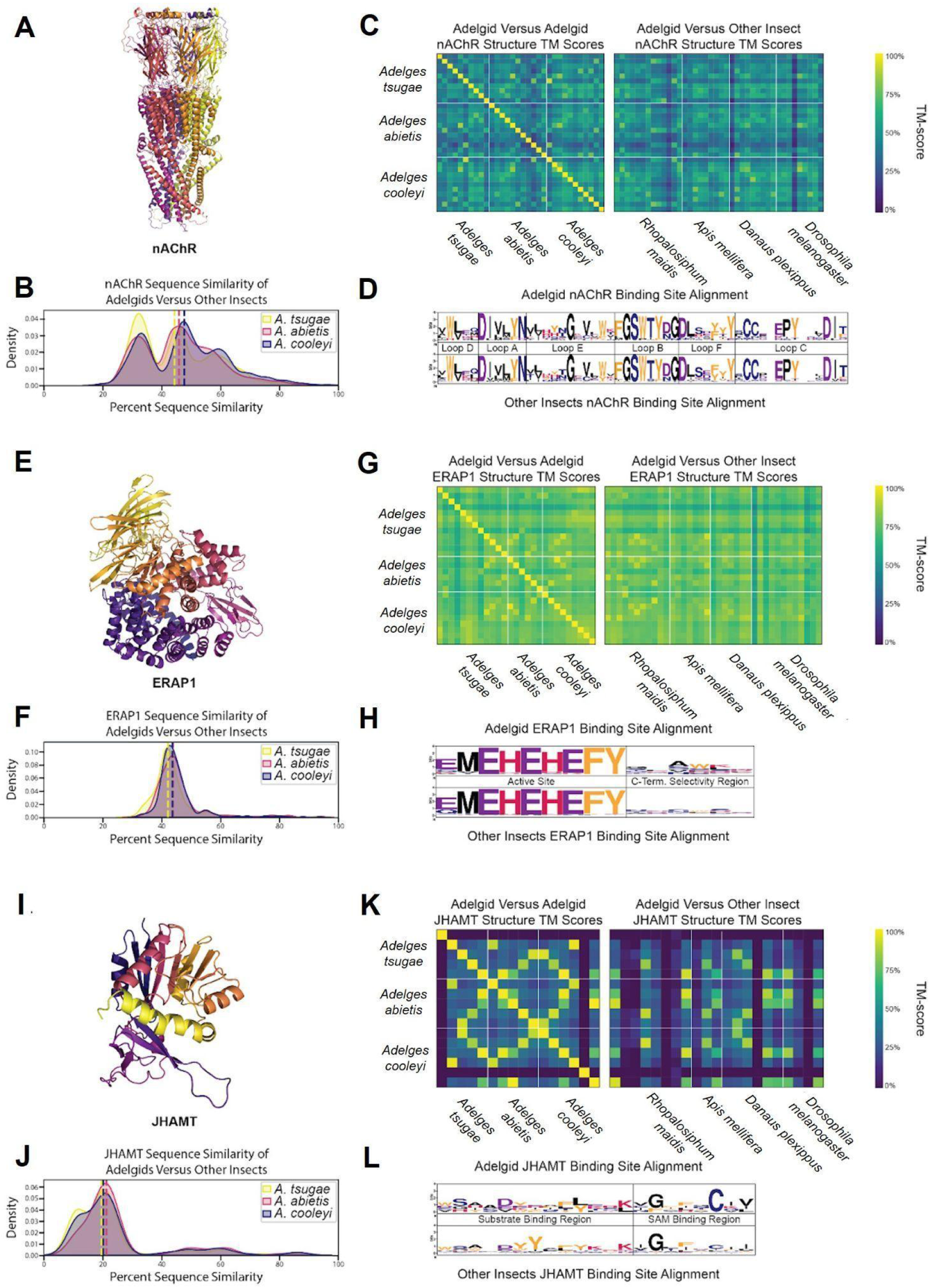
(A) AlphaFold 3 structure of an *A*. *tsugae* nAChR. (B) KDE plot of *A. tsugae*, *A. abietis*, and *A. cooleyi* percent sequence similarity compared to other insect nAChR proteins. (C) Structure similarity heatmap comparing adelgid and non-adelgid nAChR proteins. (D) Logo plot of the consensus sequences of nAChR ligand binding sites, separated into active site residues and C-terminal selectivity region residues. (E) AlphaFold 3 structure of an *Adelges tsugae* ERAP1. (F) Kernel density estimate (KDE) plot of *A. tsugae, A. abietis,* and *A. cooleyi* percent sequence similarity compared to other insect (*R*. *maidis*, *A*. *mellifera*, *D*. *plexippus*, and *D. melanogaster*) for ERAP1 proteins. (G) Structure similarity heatmap comparing adelgid and non-adelgid ERAP1 proteins. The number of proteins per heat map varies with the copy number. (H) Logo plot of the consensus sequences of ERAP1 ligand binding sites, labeled by binding loop identity. (I) AlphaFold 3 structure of an *A*. *tsugae* JHAMT. (J) KDE plot of *A. tsugae*, *A. abietis*, and *A. cooleyi* percent sequence similarity compared to other insect JHAMT proteins. (K) Structure similarity heatmap comparing adelgid and non-adelgid JHAMT proteins. (L) Logo plot of the consensus sequences of JHAMT ligand binding sites, separated into substrate binding site and S-Adenosyl methionine (SAM) binding site.

Many of the most commonly used chemical insecticides target insect nAChRs (Millar and Denholm 2007). One such insecticide, imidacloprid, is currently the standard of care for trees infested with *Adelges tsugae*, but its use is complicated by its broad-spectrum action against non-target insects (Eisenback et al. 2010). A better understanding of nAChR structural diversity could potentially help guide the discovery of precision insecticides that avoid these off target effects. Adelgid nAChR show moderate conservation, with an average of 46% across different insect taxa, and with occasional higher similarity pairs (>70%) that likely indicate nAChRs of the same or similar subtypes, and some low similarity pairs as outliers (Figure 4B). The 3D structural similarity also follows a moderate to high pattern (>0.65), with scores more evenly distributed and no notable outliers (Figure 4C). nAChR ligand binding sites, distinguished by loop identity, were highly conserved across all species studied, displaying minimal variability in key binding residues (Figure 4D). Overall, predicted adelgid nAChR structures generally align well with known human crystal structures, indicating a high level of conservation which extends across animals and indicates a vital conservation of function (PDB ID: 5FJV, Figure S1). While this conservation confirms nAChRs as functionally critical targets for pest control due to their essential role in neural signaling, the structural similarity to human nAChRs underscores the importance of developing precision insecticides that exploit adelgid-specific features to minimize off-target effects in non-target organisms.

### Effector gene families

Hemipterans use effector proteins to manipulate host plant defenses for successful feeding (Huiying Wang, Shi, and Hua 2023; Kaloshian and Walling 2016; Ray and Casteel 2022), and adelgids, as members of Hemiptera, likely employ similar strategies. Analysis of effector gene families revealed significant expansions in *Adelges tsugae* and *A. cooleyi*, particularly in *Endoplasmic reticulum aminopeptidase* 1 (*ERAP1*). ERAP proteins belong to the M1 family of zinc-dependent aminopeptidases, characterized by specific conserved structural motifs essential for their function (Figure 4E). *ERAP1* copy numbers are elevated in *Adelges tsugae* (12 copies), *A. cooleyi* (12) (Figure 2C, 2D), and *R. maidis* (14) compared to non-hemipteran insects like *A. mellifera* (8) and *D. plexippus* (10) (Table S22). This pattern mirrors findings in *Acyrthosiphon pisum*, where *ERAP1*-like C-terminal domains were enriched in salivary effectors, implicating these proteins in disrupting plant immune signaling through peptide hydrolysis of host defense molecules (Boulain et al. 2018; Chen et al. 2019).

In order to better understand the diversity of ERAP1 structures across insects, structural analysis was conducted. ERAP1 exhibited low interspecies variability and sequence similarity was consistently moderate, with an average value of 43% (Figure 4F). By translating the ERAP1 sequences into predicted structures in AlphaFold3, high structural similarity was observed, with an average TM-score of 0.87, indicating that its overall 3D structure is conserved (Figure 4G). The catalytic site of ERAP1 was also highly conserved across all species (Figure 4H). In contrast, the region of the binding pocket known for substrate C-terminal binding, which influences substrate selectivity, was highly variable. This suggests that while the overall structures are largely similar, substrate selectivity among these proteins may differ (Giastas et al. 2019). The overall predicted structures of insect ERAP1 overlapped remarkably well with human ERAP1 crystal structures, further indicating a high level of structural conservation among eukaryotes (PDB ID: 6RQX, Figure S2).

### Developmental gene families show potential for selective adelgicide development

Juvenile hormones (JH), critical sesquiterpenoid regulators of development and reproduction in insects (Shinoda and Itoyama 2003), show contrasting evolutionary trajectories in *Adelges tsugae* gene families. While *juvenile hormone acid methyltransferase* (*JHAMT*), a ∼33 kDa enzyme catalyzing the final JH biosynthesis step via SAM-dependent methylation of JH acids (Figure 4I) underwent contraction in *A. tsugae* (5 copies). In contrast, haemolymph juvenile hormone binding protein (JHBP) expanded markedly in *A. tsugae* (9 copies). This expansion pattern is consistent with *A. cooleyi* (*JHAMT*: 6 copies; *JHBP*: 9 copies) but contrasts sharply with *A. abietis*, which retained ancestral JHAMT levels (5 copies), with JHBP expanded to 20 copies (Table S21).

JHAMT has been implicated as a viable target for insect control, with RNAi feeding studies disrupting development in *B*. *dorsalis (Zhou et al. 2022)*. As JH signaling is generally unique to insects, JHAMT represents an opportunity for the development of chemical insecticides with little risk of off-target effects. JHAMT was the least conserved among the three proteins studied, with an average sequence similarity of 20% compared to the other adelgids and insects (Figure 4J). Within adelgids, JHAMT proteins showed high similarity, but only low to moderate similarity when compared to the other insects. Structural similarity mirrored this pattern, displaying occasional high-scoring matches but generally exhibiting considerable structural diversity (Figure 4K). The substrate binding region was also quite variable, with known binding residues from previous studies aligning poorly with the sequences observed in adelgids (Figure 4L) (Guo et al. 2021). The predicted structures of adelgid JHAMT enzymes differ greatly from known crystal structures of JHAMT, suggesting again that these enzyme’s structures are highly variable (PDB ID: 7EC0, Figure S1). To explore the potential of JHAMT as a selective adelgicide target, molecular docking experiments were applied to the homology model of JHAMT. This is a standard approach in modern computational drug discovery (Sadybekov and Katritch 2023), but its application here to forest health is a novel application of conservation chemistry principles. The overlapping JHAMT homology protein model from six insect species generated by AlphaFold 3 showed the amino acid differences in the binding pocket of JHAMT in *A. tsugae* (Figure 5A). The docking scores of JHAMT inhibitors (3-deazaneplanocin A, JHSI48) and ligand (*S*-adenosyl-L-homocysteine, SAH) against six JHAMT proteins showed that these ligands bound less favorably to the *A. tsugae* pocket than to that of other insects (Figure 5B). A pesticide library and the DrugBank library (Knox et al. 2024) were used for the virtual screening experiment to identify molecules with better protein-ligand interactions compared to the reported inhibitors. An analysis of docking scores for both virtual screening experiments showed that JHAMT from all insects had a diversity of interactions with the ligands (Figure 5C). A survey of the docking results however revealed that zelandopam, a dopamine-D1 receptor agonist which was developed as a hypertension medicine for humans, was found to interact selectively with *A. tsugae’s* JHAMT binding pocket, with a docking score of –10.4 kcal/mol compared to –6.9 kcal/mol for *Apis mellifera* (Figure 5B). These results support the potential for discovering selective insecticides for conservation purposes (Juang and Cernak 2025)

**Figure 5.**
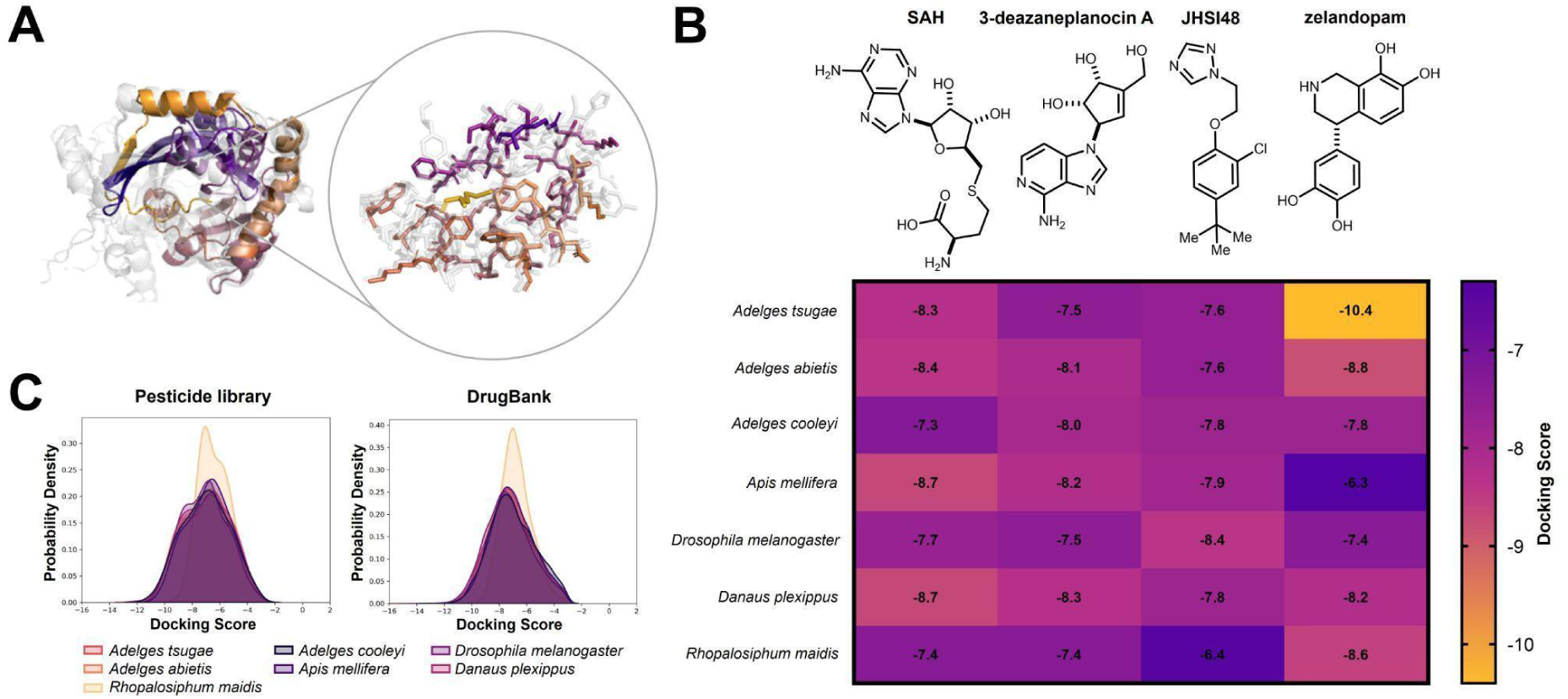
Computational analysis of JHAMT homology protein models reveals the potential for selective targeting strategy of *A. tsugae* over other insects. The protein homology models were generated by Alphafold 3.(A) *A. tsugae* JHAMT binding pocket overlaid with six other insect species highlighting the binding site amino acids in *A. tsugae*. Colored: *A. tsugae*; Gray: six other insect species. (B) Computational docking scores of the JHAMT substrate SAH, known JHAMT inhibitors (3-deazaneplanocin A and JHSI48), and DrugBank compound (zelandopam). Lower docking scores predict stronger protein-ligand interactions. SAH: S-adenosyl-L-homocysteine. (C) Kernel density estimate analysis of computed docking scores using a pesticide library and DrugBank as ligands for JHAMT.

Another rapidly expanding gene famliy in *A. tsugae* was *Survival of Motor Neuron* (*SMN*). SMN proteins are evolutionary conserved components of the spliceosomal machinery (Sm), essential for small nuclear ribonucleoprotein (snRNP) assembly and RNA processing (Chaytow et al. 2018). They contain three domains: Gemin2-binding complex which facilitates binding to gemin proteins required for the snRNP assembly (Singh et al. 2017), Tudor domain which binds symmetrically dimethylated arginine residues in Sm proteins (Selenko et al. 2001); and YG-Box domain that mediates oligomerization of *SMN* subunits, essential for functional *SMN* complexes (Praveen et al. 2014). *SMN* orchestrates snRNP biogenesis by chaperoning Sm proteins to spliceosomal RNAs, a depletion of which causes impairment to neuronal development in insects like *D. melanogaster* (Grice and Liu 2011). The *SMN* family was expanding in *A. tsugae* (1 copy), neutral in *A. cooleyi* (3 copies), and contracting in *A. abietis* (no copies), whereas *D. plexippus* had 2 copies, and *D. melanogaster* had one copy (Table S22).

SMN was deprioritized as a target for several key reasons. First, the inconsistent evolutionary patterns across adelgid species suggest that SMN function may not be universally critical for adelgid survival – *A. abietis* appears to have completely lost SMN copies yet remains a successful pest, indicating functional redundancy or alternative pathways. Second, the fundamental role of SMN in RNA processing makes it a poor candidate for selective targeting, as disrupting such basic cellular machinery would likely affect all eukaryotic organisms rather than providing adelgid-specific vulnerability. Finally, the variable copy numbers across closely related adelgid species (ranging from 0 to 3 copies) suggest that targeting SMN would not provide consistent efficacy across different adelgid populations or species.

## Conclusion

The high-quality genome assemblies presented here for *Adelges tsugae* and *A. abietis* reveal conservation in genome architecture, from chromosome-level synteny to shared patterns of repeat evolution and gene family dynamics. Gene family expansions in *A. abietis* included metabolic gene families, involved in carbohydrate transport, while expansions in *A. tsugae* represented genes involved in cell differentiation and development, and suppression of plant immunity. Genes associated with nicotinic acetylcholine receptors (*nAChR*s) exhibited strong conservation across species, reflecting their conserved role in neural signaling. The *Endoplasmic reticulum aminopeptidase 1* (*ERAP1*) gene family exhibited expansion, and *juvenile hormone acid methyltransferase* (*JHAMT*) displayed contraction relative to ancestral states. While nAChR and ERAP1 had similar three-dimensional protein structures across insects tested, JHAMT exhibited notable variability in key regions, pointing to functional divergence. The opportunity to potentially disrupt juvenile hormone signaling pathways for adelgid-selective pest control was explored by virtual drug and pesticide screening, which identified the hypertension drug zelandopam as having selective interactions with JHAMT in hemlock woolly adelgid.

## Supporting information

Supplemental Files ReadMe

## Acknowledgments

The authors acknowledge the contributions of the HPC resources from the Computational Biology Core and support for sequencing through the Center for Genome Innovation, both within the Institute for Systems Genomics at the University of Connecticut. We would also like to thank Dr. Gaelen Burke and Dr. Dustin Dial at the University of Georgia for providing laboratory space as well as Dr. Ben Smith at the Mountain Research Station in North Carolina. Dr. Nathan Havill of the USDA Forest Service and Dr. Bryan Brunet of Agriculture and Agri-Food Canada are thanked for providing preliminary nAChR sequences.

## Data availability

All scripts and data are described in https://gitlab.com/PlantGenomicsLab/adelges-tsugae-genomics. NCBI BioProject ID PRJNA1163707 contains the raw sequencing reads for *Adelges tsugae* and *A. abietis*. RNA reads (SRR30936311 ONT long reads for *A. tsugae*, SRR31285667 HiC reads for *A. abietis*, SRR31285820-3 10x linked reads for *A. abietis*, SRR30936309 and SRR30936310 short RNA reads for *A. tsugae*). The genome assemblies, structural and functional annotations are also hosted in the Gitlab.

