## Supplemental Figure 1 for "Chromosome scale genomes of two invasive Adelges species enable virtual screening for selective adelgicides"

**Supplemental_Figure_S1:** Hi-C contact map analysis and manual curation of *Adelges abietis*. (A) Hi-C contact map before manual curation, showing structural inconsistencies in Chromosome 1. (B) Zoomed-in view of the contact map reveals evidence of heterozygous reciprocal translocations, suggesting two haplotype configurations: {AB} {CD} and {AD} {CB}. These patterns indicate a potential fusion between *A. cooleyi* Chr2 and a large portion of Chr10. (C) Manual curation addressed these inconsistencies by splitting Chromosome 1 at the identified breakpoint based on Hi-C signal patterns, resulting in two separate chromosomes to better reflect the true chromosomal structure. (D) Zoomed-in view of the split Chromosome 1, showing scaffold lengths and improved resolution after curation.

A

B

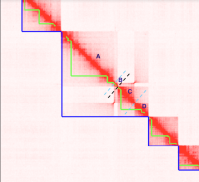

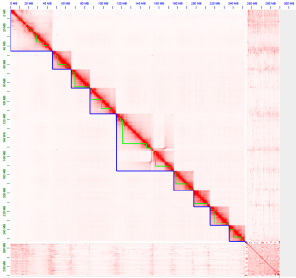

C	D

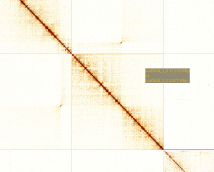

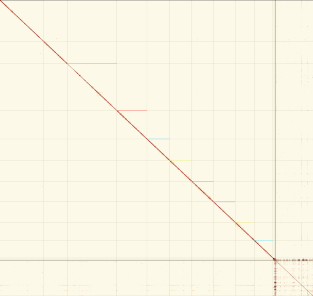
