## Supplemental Files ReadMe for "Chromosome scale genomes of two invasive Adelges species enable virtual screening for selective adelgicides"

**Supplemental_Tables_S1-S20.xlsx**

**Supplemental_Table_S1:** RNA read statistics for *A. tsugae* and *A. abietis*.

**Supplemental_Table_S2:** Completeness and statistics of genome annotations across Insecta.

**Supplemental_Table_S3:** Gene orthogroup file from all annotations of 45 species across Hemiptera. Rooted phylogeny ultrametric tree:

(Tetrancychus_urticae:1,(Daphnia_pulex:0.639,((((Apis_mellifera:0.290877,Nasonia_vitripennis:0.290877):0.220279,(((Drosophila_melanogaster:0.357663,Anopheles_gambiae:0.357663):0.0914771,(Danaus_plexippus:0.169979,(Bombyx_mori:0.142965,Chilo_suppressalis:0.142965):0.0270142):0.279162):0.0293862,(Anolophora_glabripennis:0.244311,Tribolium_castaneum:0.244311):0.234216):0.0326291):0.0255772,(Pediculus_humanus:0.504826,((Thrips_palmi:0.17608,Frankliniella_occidentalis:0.17608):0.283375,Zootermopsis_nevadensis:0.459455):0.0453713):0.0319067):0.0326411,(((Bemisia_tabaci:0.475285,(Diaphorina_citri:0.250355,Pachypsylla_venusta:0.250355):0.22493):0.0313343,((Parthenolecanium_corni:0.283013,Planococcus_citri:0.283013):0.186725,((Daktulosphaira_vitifoliae:0.13439,(Adelges_tsugae:0.0575452,(Adelges_abietis:0.0450085,Adelges_cooleyi:0.0450085):0.0125368):0.0768448):0.0595904,(Cinara_cedri:0.148596,(Sipha_flava:0.119623,((Diuraphis_noxia:0.0517579,((Metopolophium_dirhodum:0.0162814,Acyrthosiphon_pisum:0.0162814):0.0190782,Myzus_persicae:0.0353597):0.0163982):0.017431,(((Rhopalosiphum_maidis:0.0118807,Rhopalosiphum_padi:0.0118807):0.0363513,((Aphis_glycines:0.0259716,Aphis_gossypii:0.0259716):0.00899398,Aphis_craccivora:0.0349656):0.0132665):0.00518617,Melanaphis_sacchari:0.0534182):0.0157707):0.0504344):0.0289727):0.0453844):0.275758):0.0368816):0.025254,(((Laodelphax_striatellus:0.156051,Nilaparvata_lugens:0.156051):0.290768,(Macrosteles_quadrilineatus:0.229654,Homalodisca_vitripennis:0.229654):0.217166):0.0482858,((Cimex_lectularius:0.319189,(Nesidiocoris_tenuis:0.196228,Apolygus_lucorum:0.196228):0.122961):0.0571672,(Nezara_viridula:0.0861208,Halyomorpha_halys:0.0861208):0.290235):0.118749):0.0367687):0.0375002):0.0696263):0.361):0

Unrooted phylogeny ultrametric tree with support values:

(Tetrancychus_urticae:0.627297,Daphnia_pulex:0.462404,((((((Drosophila_melanogaster:0.381062,Anopheles_gambiae:0.350536)0.856209:0.100728,(Danaus_plexippus:0.13056,(Chilo_suppressalis:0.145604,Bombyx_mori:0.131594)0.461538:0.0279979)0.9819:0.258301)0.329311:0.0324811,(Tribolium_castaneum:0.163897,Anolophora_glabripennis:0.175947)0.956762:0.152982)0.244344:0.0304611,(Nasonia_vitripennis:0.209478,Apis_mellifera:0.197757)0.95827:0.148127)0.128205:0.0213031,(Pediculus_humanus:0.341026,((Thrips_palmi:0.123655,Frankliniella_occidentalis:0.132293)0.964304:0.211382,Zootermopsis_nevadensis:0.256124)0.268477:0.0288937)0.0995475:0.0203666)0.0648567:0.0244061,((((Diaphorina_citri:0.203929,Pachypsylla_venusta:0.226182)0.8909:0.203727,Bemisia_tabaci:0.316329)0.135747:0.025029,((((Sipha_flava:0.0839266,((((Rhopalosiphum_padi:0.00642295,Rhopalosiphum_maidis:0.00774381)0.794872:0.0146569,((Aphis_gossypii:0.00634676,Aphis_glycines:0.0494299)0.531926:0.0117771,Aphis_craccivora:0.035962)0.618904:0.0199239)0.251885:0.00567592,Melanaphis_sacchari:0.0271924)0.468577:0.0127376,((Myzus_persicae:0.0199007,(Metopolophium_dirhodum:0.0100843,Acyrthosiphon_pisum:0.0143654)0.751634:0.0135529)0.410256:0.00798164,Diuraphis_noxia:0.0558148)0.507793:0.0133056)0.674208:0.0389595)0.403218:0.0196119,Cinara_cedri:0.165739)0.61991:0.0434209,(Daktulosphaira_vitifoliae:0.0990772,(Adelges_tsugae:0.0440353,(Adelges_cooleyi:0.0201619,Adelges_abietis:0.0579179)0.473102:0.0113754)0.928607:0.0653067)0.802916:0.0459908)0.918049:0.248797,(Parthenolecanium_corni:0.258687,Planococcus_citri:0.214656)0.928105:0.159863)0.311212:0.0342)0.12368:0.0232983,(((Nezara_viridula:0.068875,Halyomorpha_halys:0.0495769)0.951735:0.196166,((Nesidiocoris_tenuis:0.155453,Apolygus_lucorum:0.162045)0.898441:0.105274,Cimex_lectularius:0.209838)0.578683:0.0445369)0.806435:0.0878268,((Nilaparvata_lugens:0.108384,Laodelphax_striatellus

**Supplemental_Table_ S4:** Full amino acid sequences of proteins used for heat map and logo plot analyses.

**Supplemental_Table_S5:** Residue-by-residue alignments used to create ERAP1 logo plot for Figure 4D.

**Supplemental_Table_S6:** Residue-by-residue alignments used to create JHAMT logo plot for Figure 4H.

**Supplemental_Table_S7:** Residue-by-residue alignments used to create nAChR logo plot for Figure 4L.

**Supplemental_Table_S8:** Individual logo plot sequences for each studied species.

**Supplemental_Table_S9:** DNA read statistics for *Adelges tsugae*.

**Supplemental_Table_S10:** Initial genome assembly statistics for *A. tsugae* and *A. abietis*. Quality assessment statistics are based on contigs of size >= 500 bp, unless otherwise noted (e.g., "# contigs (>= 0 bp)" and "Total length (>= 0 bp)" include all contigs). Dependencies and versions for completeness estimates: hmmsearch: 3.1, bbtools: 39.01, metaeuk: 6.a5d39d9, busco: 5.4.5.

**Supplemental_Table_S11:** Read-level DNA contaminant filtering results for *A. tsugae*.

**Supplemental_Table_S12:** Contig-level DNA contaminant filtering results for *A. tsugae* and *A. abietis* initial assemblies.

**Supplemental_Table_S13:** Repeat statistics for *A. tsugae* and *A. abietis.*

**Supplemental_Table_S14:** Structural annotation statistics for *A. tsugae* and *A. abietis.*

**Supplemental_Table_S15:** Functional annotation statistics for *A. tsugae* and *A. abietis*.

**Supplemental_Table_S16:** Synteny analysis between *Daktulosphaira vitifoliae*, *Adelges cooleyi* (top 10 scaffolds), *A. abietis*, and *A. tsugae*.

**Supplemental_Table_ S17:** Gene density of chromosome X1 on *A. abietis* and *A. tsugae*

**Supplemental_Table_ S18:** Gene orthogroup description file showing only the 244 significantly divergent orthogroups before filtering transposable elements.

**Supplemental_Table_S19:** Gene orthogroup description file showing only the 235 significantly divergent orthogroups after filtering transposable elements.

**Supplemental_Table_S20:** Phylogenetic analysis of all orthogroups across Hemipterans.

**Supplemental_Table_S21:** Lineage-specific rapidly evolving HOGs in *A. tsugae, A. abietis*, *A. cooleyi,* the most recent common ancestry (MRCA) of *Adelges*, and the MRCA of *A. cooleyi* and *A. abietis*.

**Supplemental_Table_S22:** Copy numbers and gene identifiers of the ERAP, SMN, JHBP, and JHAMT gene families in Insecta.

**Supplemental Files**

**Supplemental_File_S1:** Full Protocol with Modifications "Manual Purification of High-MolecularWeight Genomic DNA from Fresh or Frozen Tissue."

**Supplemental_File_S2:** Final basecalling report for *Adelges tsugae* DNA reads.

**Supplemental Figures**

**Supplemental_Figure_S1:** Hi-C contact map analysis and manual curation of *A. abietis*. (A) Hi-C contact map before manual curation, showing structural inconsistencies in Chromosome 1. (B) Zoomed-in view of the contact map reveals evidence of heterozygous reciprocal translocations, suggesting two haplotype configurations: {AB} {CD} and {AD} {CB}. These patterns indicate a potential fusion between *A. cooleyi* Chr2 and a large portion of Chr10. (C) Manual curation addressed these inconsistencies by splitting Chromosome 1 at the identified breakpoint based on Hi-C signal patterns, resulting in two separate chromosomes to better reflect the true chromosomal structure. (D) Zoomed-in view of the split Chromosome 1, showing scaffold lengths and improved resolution after curation.

**Supplemental_Figure_S2:** Three-dimensional protein models for ERAP, JHAMT, and nAChR overall and active site models, with overlaid crystal structures.
