## Supplemental Figure 2 for "Chromosome scale genomes of two invasive Adelges species enable virtual screening for selective adelgicides"

**Supplemental_Figure_S2:** Three-dimensional protein models for ERAP, JHAMT, and nAChR overall and active site models, with overlaid crystal structures.

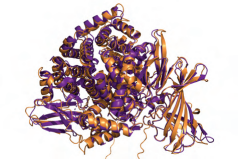

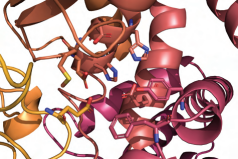

ERAP1 AF3 Model Overlaid With Crystal Structure (PDB ID: 6RQX)

ERAP1 AF3 Model Active Site

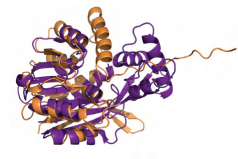

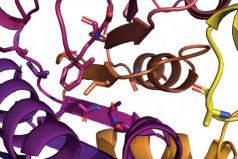

JHAMT AF3 Model Overlaid With Crystal Structure (PDB ID: 7EC0)

JHAMT AF3 Model Active Site

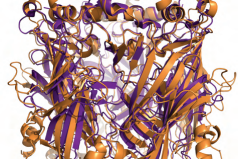

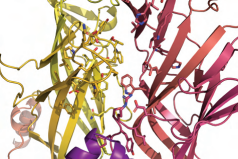

nAChR AF3 Model Overlaid With Crystal Structure (PDB ID: 5FJV)

nAChR AF3 Model Active Site
