## Supplemental File 1 for "Chromosome scale genomes of two invasive Adelges species enable virtual screening for selective adelgicides"

**Supplemental_File_S1:** Full Protocol with Modifications "Manual Purification of High-MolecularWeight Genomic DNA from Fresh or Frozen Tissue."

*Adelges tsugae* Extraction protocol modified from QIAGEN MagAttract® HMW DNA Handbook published 03/2020:

Fresh and frozen tissue can be used for DNA purification. Yield and the DNA quality obtained will depend on the tissue type, source, and storage conditions.

Important points before starting:

- To ensure that the extracted DNA is not fragmented, avoid shearing stress such as fast or unnecessary pipetting steps.

- Ensure that Buffers MW1 and PE have been prepared according to the instructions. - Ensure that the magnetic particles are fully resuspended. Vortex the vessel containing the magnetic particles vigorously for at least 3 min before first use.

Procedure:

1. Place insect samples (single or multiple) in a 1.5 ml tube and set on ice. 2. Crush insects with a pestle in the small liquid nitrogen dewer until it is a fine powder. 3. Add 200ul T1 lysis buffer to the tube and gently mix (it can be frozen at -80oC at this point if desired). 4. Add 20 μl Proteinase K. Mix by inverting the tube 5 times.

5. Add 4 μl RNAse A and 150 μL T1 Buffer to the sample. Mix by inverting the tube 5 times. 6. Incubate 20 hours at 32°C at 5 rpm in Illumina Hybridization Oven. Add 4 μL RNase A. *Modified from: “Incubate the sample for 2 hours at 27°C and 600 rpm.”*

7. Centrifuge briefly.

8. Transfer 350-370 µl of the lysate to a new 2 ml sample tube.

*Note*: We do not recommend the use of 1.5 sample tubes because of the increased risk of salt carryover into the eluate due to inefficient mixing caused by the conical shape of the tube.

*Note*: If pieces of insoluble material are still present, centrifuge at 20,000 x g for 3 min and transfer the supernatant into a clean sample tube.

9. Vortex the MagAttract® Suspension G for 1 min and add 15 μl to the sample. *Note*: If this is the first time using MagAttract Suspension G, increase the vortexing time to 3 min. 10. Add 280 μl Buffer MB. Incubate at room temperature for 3 min at 1400 rpm. 11. Centrifuge the tube briefly and place on the MagAttract Magnetic Rack, wait until bead separation has been completed (~1 min), and remove and discard the supernatant. *Note*: Avoid disturbing the magnetic bead pellet while aspirating the supernatant. Remove the supernatant completely. 12. Add 700 µl Buffer MW1 to the sample. Incubate at room temperature for 1 min at 1400

rpm. *Note*: To increase the efficiency of the wash step, remove the tube from the MagAttract Magnetic Rack before adding the wash buffer. Add the wash buffer directly onto the magnetic bead pellet.

13. Centrifuge the tube briefly and place on the MagAttract Magnetic Rack, wait until bead separation has been completed (~1 min), and remove and discard the supernatant. 14. Repeat steps 12 and 13 one time.

15. Add 700 µl Buffer PE to the sample. Incubate at room temperature for 1 min at 1400 rpm. *Note*: To increase the efficiency of the wash step, remove the tube from the MagAttract

Magnetic Rack before adding the wash buffer. Add the wash buffer directly onto the magnetic bead pellet.

16. Centrifuge the tube briefly and place on the MagAttract Magnetic Rack, wait until bead separation has been completed (~1 min), and remove and discard the supernatant. 17. Repeat steps 15 and 16.

*Note*: Remove all the supernatant. Use a small pipette tip to remove any traces of Buffer PE.

18. Rinse the particles with 700 µl distilled water while the tube is on the MagAttract Magnetic Rack and the beads are fixed to the wall of the sample tube. Incubate for exactly 1 min at room temperature and remove the supernatant.

Important: Do not pipet water directly onto the bead pellet – pipet it into the sample tube against the side facing away from the bead pellet. All pipetting steps must be performed carefully to avoid disturbing the fixed bead pellet.

19. Repeat step 18.

20. Remove the tube from the magnetic rack and add an appropriate volume of Buffer AE (100– 200 µl). Incubate at room temperature for 3 min at 1400 rpm.

21. Centrifuge the tube briefly and place the tube on the MagAttract Magnetic Rack, wait until bead separation has been completed (~1 min), and transfer the supernatant with the high molecular weight DNA to a new sample tube using a wide-bore pipette tip.

*Note*: The yield of genomic DNA depends on the sample type and the number of cells in the sample. For some downstream applications, concentrated DNA may be required. Elution with volumes <200 µl (e.g., 150 µl or 100 µl) increases the DNA concentration in the eluate.

22. Elute with Buffer AE at 37°C, 3 minutes at 1400rpm then 7 minutes stationary *Modified from: “A second elution with Buffer AE will increase the total DNA yield. Due to the increased volume, the DNA concentration is reduced.”*

23. Store the extracted gDNA sample at 4°C for up to 2 weeks or at −20°C for up to 6 months or proceed directly to GEM Generation & Barcoding.

Protocol for *Adelges abietis*:

Genomic DNA:

The *Adelges abietes* genome sequencing library was prepared using the 10X Genomics Linked Reads kit. Hi-C libraries were prepared and sequenced by Phase Genomics. Approximately 30 *Adelges abietis* individuals were gently crushed with a plastic pestle in a 1.5mL Eppendorf tube containing 600 microliters Lysis Buffer (10 mM Tris pH 7.5, 400 mM NaCl, 100 mM EDTA), 40 microliters 10% Sodium dodecyl sulfate, and 100 microliters Proteinase-K solution (20 mg/mL). Sample was incubated at 37°C for 12 hours. After incubation, 240 microliters of 5M NaCl were added and the reaction was mixed by inverting the tube five times. Sample was centrifuged for 15 minutes at 1100 x g at 4°C and supernatant was transferred to a new 1.5 mL Eppendorf tube. 1.2 mL of absolute ethanol was added and the sample was centrifuged at 6250 x g for 5 minutes at 4°C. The supernatant was removed and the sample was air dried for five minutes. 35 microliters of TE (10 mM Tris pH 8.0, 0.1 mM EDTA) was added to the pellet.

Linked-read DNA: The single *Adelges abietis* was frozen in liquid nitrogen in a plastic Eppendorf tube and ground using a plastic pestle. One mL 1% Formaldehyde was added and the sample was placed on a nutator for 20 min at room temperature. 110 uL of 1.25M Glycine was added and the sample was placed on a nutator for 15 min at room temperature. The fixed ground tissue was spun down in a tabletop microcentrifuge at 15,000 X g for 2 min. Most supernatant was gently removed with a pipeteman and replaced with 500 uL Phosphate Buffered Saline (PBS). The sample was spun a second time under the same conditions, most of the supernatant was removed, and the sample was stored at -80°C.
