## Supplemental File 2 for "Chromosome scale genomes of two invasive Adelges species enable virtual screening for selective adelgicides"

**Supplemental_File_S2:** Final basecalling report for *Adelges tsugae* DNA reads.

**PromethION 24 (PCA100015) Final report**

Dec 18, 23, 3:16 PM UTC-4:00 — Dec 22, 23, 3:18 PM UTC-4:00 ·

2023DEC18_Wegrzyn_HWA_fulllength_clean · 2023DEC18_Wegrzyn_HWA_fulllength_clean · 3C Protocol run ID: 1b7971fb-bf36-402e-a5a1-030b814af096

Run summary Run configuration Sequence output Run health Run log **Run summary**

**DATA OUTPUT**

Estimated bases

**100.46 Gb**

Estimated N50

**10.5 kb**

**BASECALLING**

Reads called

**100%**

**Run configuration RUN SETUP**

Reads generated

**32.5 M**

Total data produced (pass / fail) **1.33 TB**

Bases called (min Q score: 10) **89.94 Gb 10.84 Gb** Pass Fail

**RUN DURATION**

Run time

96 hrs 0 mins / 96 hrs 0 mins (est)

Elapsed time Run limit

Run status

***FINISHED*** *· Target runtime has been reached*

View unit abbreviations used in this report

**DATA OUTPUT SETTINGS**

Flow cell type FLO-PRO114M Flow cell type alias FLO-PRO114M Flow cell ID PAU00939

Kit type SQK-LSK114

**RUN SETTINGS**

Run limit 96 hrs

Active channel selection On

Pore scan freq. 1.5 hrs

Reserved pores On

Minimum read length 200 bp

Read splitting On

Basecalling Super-accurate basecalling, 400 bps

Modified basecalling On

Modified base context 5mC & 5hmC **Sequence output**

FAST5 output Off

FASTQ output gzip_compress

FASTQ reads per file 4000

BAM output On

Bulk file output Off

Data location /data/./2023DEC18_Wegrzyn_ HWA_fulllength_clean/2023DE

C18_Wegrzyn_HWA_fulllength

_clean/20231218_1516_3C_P

AU00939_1b7971fb

**SOFTWARE VERSIONS**

MinKNOW 23.07.12

Bream 7.7.6

Configuration 5.7.11

Guppy 7.1.4

MinKNOW Core 5.7.5

**READ LENGTHS · OUTLIERS REMOVED OUTLIERS**

The read length graph shows the total number of bases vs the read length. The longest 1% of strands are classified as outliers, and excluded to allow focus on the main body of data.

The longest 1% of strands are classified as outliers, and aggregated into groups to show

Legend

Basecalled Estimated

Estimated N50 **10.5 kb**

% Basecalled **100%**

their relative amounts.

Read

length (kb)

Aggregated reads (Mb)

80 - 144 849.24
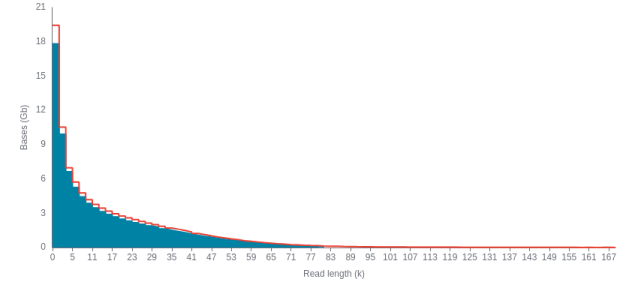

144 - 208 48.55

208 - 260 1.63

**CUMULATIVE OUTPUT**

The cumulative output shows the total amount of bases or reads sequenced over time by your device.

Bases

Legend

Estimated

Predicted total number of bases, prior to

basecalling

Passed

Bases equal to or

above the quality score threshold.

Failed

Bases below the quality score threshold.

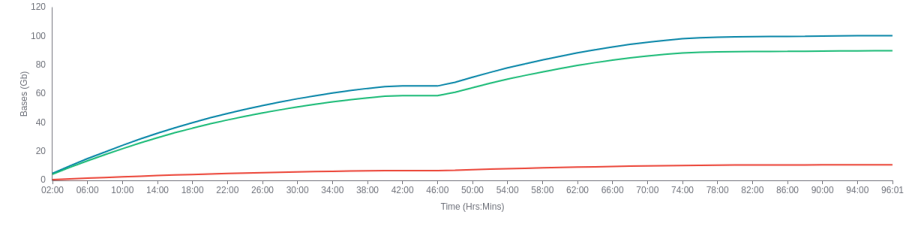
Reads

Legend

Total

Total number of reads, including passed, failed and skipped.

Passed

Reads equal to or above the quality score threshold.

Failed

Reads below the quality score threshold.

Skipped

Reads that will not be basecalled. Post run basecalling is possible.

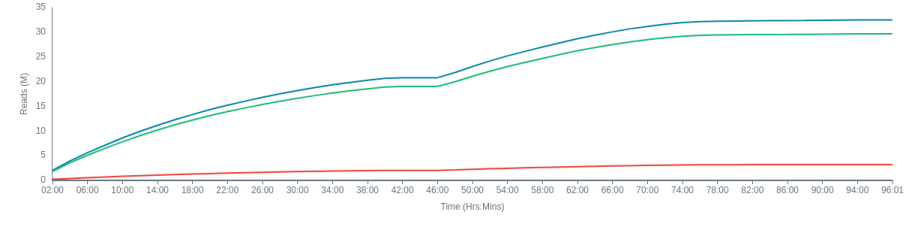

**QUALITY SCORE**

The quality score is calculated as basecalling is performed on your device. Reads that fall below the minimum value of 10 will be classified as failed reads. You can alter the accepted minimum quality score in MinKNOW.

Legend

Mode

Spread

Min. quality

The most frequent quality score of reads in the run.

The spread of quality scores, found by calculating full width half maximum.

score

Minimum quality score to be accepted as a passed read.

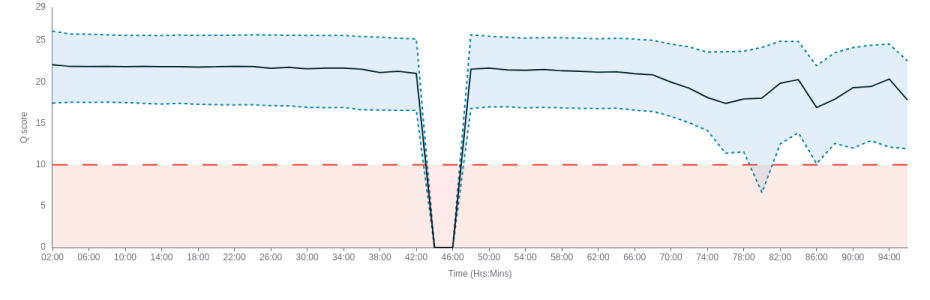

Troubleshooting

Quality score low

This can be due to the translocation speed being out of the accepted range,

which can correlate to low quality scores. If you see that the translocation speed

is out of the accepted range in the below graph, please see the Flow Cell

refuelling page linked here for further troubleshooting.

**Run health**

**PORE ACTIVITY**

The Pore activity graph shows the performance of your sample as it is being sequenced during a run.

Show grouped

Legend

Sequencing

Pore currently sequencing

Pore available

Pore available for sequencing

Unavailable

Pore currently unavailable for sequencing

Inactive

Pore no longer suitable for further sequencing

Unclassified

Pore status unknown

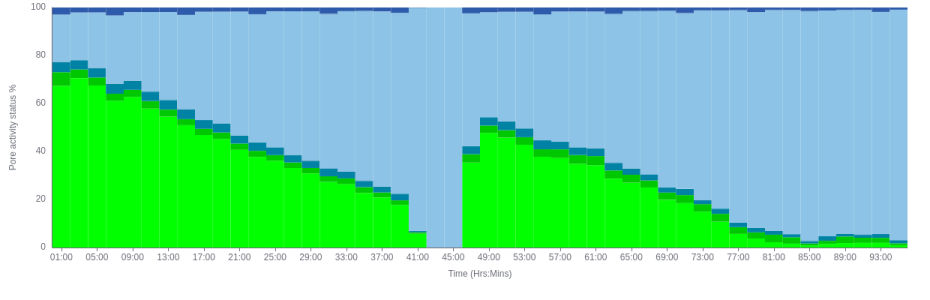

Troubleshooting

General

Some commonly seen issues are excess pores classified as Recovering, Open

Pore, or Free Adapter. To find out what advice is applicable for your run, visit the

user guide.

**PORE SCAN**

A Pore scan is performed at configurable time intervals to determine the current status of pores within channels on a Flow Cell. For this run a Pore scan is performed every 1.5 hrs.

Legend

Pore available

Pore in channel

available for

sequencing

Reserved pore

Pore in reserve, will return to available when required

Unavailable

Pore inhibited from sequencing

Saturated

Possible contamination in the sample

Zero

No current is passing through this pore,

possibly due to bubbles on the membrane

Inactive

Pore no longer suitable for further sequencing

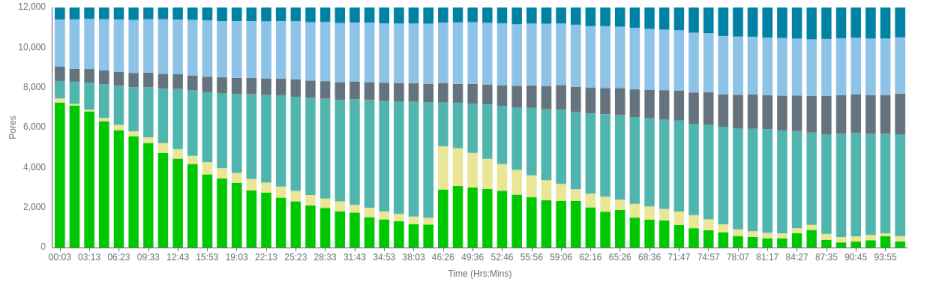
Troubleshooting

High proportion Unavailable

Possible contaminants in library blocking the pore. Consider using the Flow Cell Wash Kit, and reloading a library.

High proportion Inactive

If localised to one area of the Flow Cell, this could indicate that an air bubble has been introduced during the flushing/loading steps. If inactivity is spread across the Flow Cell this could be caused by improper loading of the library, please refer to the user guide for further support.

**TRANSLOCATION SPEED TEMPERATURE**

The translocation speed is the rate at which DNA/RNA travels through pores as it is being sequenced.

Legend

Median 75% quartile 25% quartile Accepted range

The temperature of the Flow Cell over the run time. Legend

Measured Target

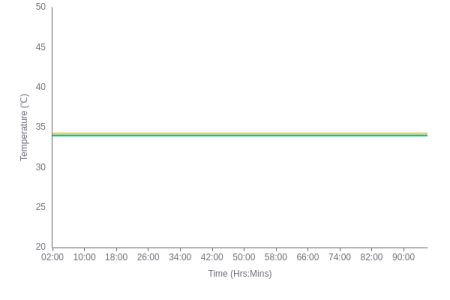

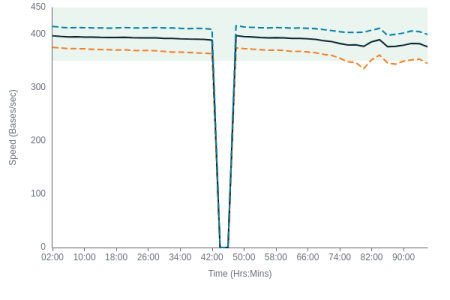
Troubleshooting

Troubleshooting

Low speed

Check that the Flow Cell is within the target temperature range.

Note

Low-quality and short reads are not included in this graph.

**Run log**

**SYSTEM MESSAGES**

Out of range

Check that the Flow Cell is correctly seated and firmly pushed down into the device. Ensure ambient temperature is always within the specified range for your device in the user guide.

Air flow should be good but not excessive. Excessive amounts of cool air blowing on the device could prevent it from reaching target temperature.

System messages are a record of the events that occurred in the time covered by this report.

**Errors**

*None*

**Warnings**

*None*

**Events**

**Disk space** · 18 Dec 23, 20:16

Disk /data has 13685 GB space remaining

**Waiting for temperature** · 18 Dec 23, 20:16

Waiting up to 300 seconds for temperature to stabilise at 34.0°C

**Starting** · 18 Dec 23, 20:18

Starting sequencing procedure

**Pore scan starting** · 18 Dec 23, 20:18

Performing Pore Scan

**Pore scan result** · 18 Dec 23, 20:21

Pore scan for flow cell PAU00939 has found a total of 7453 pores. 2517 pores available for immediate sequencing

**Message** · 18 Dec 23, 20:31

Setting temperature to reach 34.1°C

**Pore scan starting** · 18 Dec 23, 21:52

Performing Pore Scan

**Pore scan result** · 18 Dec 23, 21:56

Pore scan for flow cell PAU00939 has found a total of 7180 pores. 2505 pores available for immediate sequencing

**Pore scan starting** · 18 Dec 23, 23:27

Performing Pore Scan

**Pore scan result** · 18 Dec 23, 23:31

Pore scan for flow cell PAU00939 has found a total of 6894 pores. 2467 pores available for immediate sequencing

**Pore scan starting** · 19 Dec 23, 01:02

Performing Pore Scan

**Pore scan result** · 19 Dec 23, 01:06

Pore scan for flow cell PAU00939 has found a total of 6476 pores. 2350 pores available for immediate sequencing

**Pore scan starting** · 19 Dec 23, 02:37

Performing Pore Scan

**Pore scan result** · 19 Dec 23, 02:41

Pore scan for flow cell PAU00939 has found a total of 6138 pores. 2261 pores available for immediate sequencing

**Pore scan starting** · 19 Dec 23, 04:12

Performing Pore Scan

**Pore scan result** · 19 Dec 23, 04:16

Pore scan for flow cell PAU00939 has found a total of 5815 pores. 2265 pores available for immediate sequencing

**Pore scan starting** · 19 Dec 23, 05:47

Performing Pore Scan

**Pore scan result** · 19 Dec 23, 05:51

Pore scan for flow cell PAU00939 has found a total of 5510 pores. 2179 pores available for immediate sequencing

**Pore scan starting** · 19 Dec 23, 07:22

Performing Pore Scan

**Pore scan result** · 19 Dec 23, 07:26

Pore scan for flow cell PAU00939 has found a total of 5225 pores. 2040 pores available for immediate sequencing

**Pore scan starting** · 19 Dec 23, 08:57

Performing Pore Scan

**Pore scan result** · 19 Dec 23, 09:01

Pore scan for flow cell PAU00939 has found a total of 4923 pores. 2023 pores available for immediate sequencing

**Pore scan starting** · 19 Dec 23, 10:32

Performing Pore Scan

**Pore scan result** · 19 Dec 23, 10:36

Pore scan for flow cell PAU00939 has found a total of 4580 pores. 1975 pores available for immediate sequencing

**Pore scan starting** · 19 Dec 23, 12:07

Performing Pore Scan

**Pore scan result** · 19 Dec 23, 12:11

Pore scan for flow cell PAU00939 has found a total of 4266 pores. 1757 pores available for immediate sequencing

**Pore scan starting** · 19 Dec 23, 13:42

Performing Pore Scan

**Pore scan result** · 19 Dec 23, 13:46

Pore scan for flow cell PAU00939 has found a total of 3961 pores. 1750 pores available for immediate sequencing

**Pore scan starting** · 19 Dec 23, 15:17

Performing Pore Scan

**Pore scan result** · 19 Dec 23, 15:21

Pore scan for flow cell PAU00939 has found a total of 3731 pores. 1706 pores available for immediate sequencing

**Pore scan starting** · 19 Dec 23, 16:52

Performing Pore Scan

**Pore scan result** · 19 Dec 23, 16:56

Pore scan for flow cell PAU00939 has found a total of 3433 pores. 1529 pores available for immediate sequencing

**Pore scan starting** · 19 Dec 23, 18:27

Performing Pore Scan

**Pore scan result** · 19 Dec 23, 18:31

Pore scan for flow cell PAU00939 has found a total of 3240 pores. 1512 pores available for immediate sequencing

**Pore scan starting** · 19 Dec 23, 20:02

Performing Pore Scan

**Pore scan result** · 19 Dec 23, 20:06

Pore scan for flow cell PAU00939 has found a total of 3050 pores. 1437 pores available for immediate sequencing

**Pore scan starting** · 19 Dec 23, 21:37

Performing Pore Scan

**Pore scan result** · 19 Dec 23, 21:41

Pore scan for flow cell PAU00939 has found a total of 2828 pores. 1371 pores available for immediate sequencing

**Pore scan starting** · 19 Dec 23, 23:12

Performing Pore Scan

**Pore scan result** · 19 Dec 23, 23:16

Pore scan for flow cell PAU00939 has found a total of 2624 pores. 1288 pores available for immediate sequencing

**Pore scan starting** · 20 Dec 23, 00:47

Performing Pore Scan

**Pore scan result** · 20 Dec 23, 00:51

Pore scan for flow cell PAU00939 has found a total of 2449 pores. 1252 pores available for immediate sequencing

**Pore scan starting** · 20 Dec 23, 02:22

Performing Pore Scan

**Pore scan result** · 20 Dec 23, 02:26

Pore scan for flow cell PAU00939 has found a total of 2310 pores. 1173 pores available for immediate sequencing

**Pore scan starting** · 20 Dec 23, 03:57

Performing Pore Scan

**Pore scan result** · 20 Dec 23, 04:01

Pore scan for flow cell PAU00939 has found a total of 2124 pores. 1174 pores available for immediate sequencing

**Pore scan starting** · 20 Dec 23, 05:32

Performing Pore Scan

**Pore scan result** · 20 Dec 23, 05:36

Pore scan for flow cell PAU00939 has found a total of 1986 pores. 1010 pores available for immediate sequencing

**Pore scan starting** · 20 Dec 23, 07:07

Performing Pore Scan

**Pore scan result** · 20 Dec 23, 07:11

Pore scan for flow cell PAU00939 has found a total of 1804 pores. 982 pores available for immediate sequencing

**Pore scan starting** · 20 Dec 23, 08:42

Performing Pore Scan

**Pore scan result** · 20 Dec 23, 08:46

Pore scan for flow cell PAU00939 has found a total of 1678 pores. 935 pores available for immediate sequencing

**Pore scan starting** · 20 Dec 23, 10:17

Performing Pore Scan

**Pore scan result** · 20 Dec 23, 10:21

Pore scan for flow cell PAU00939 has found a total of 1549 pores. 835 pores available for immediate sequencing

**Pore scan starting** · 20 Dec 23, 11:52

Performing Pore Scan

**Pore scan result** · 20 Dec 23, 11:56

Pore scan for flow cell PAU00939 has found a total of 1484 pores. 855 pores available for immediate sequencing

**Pore scan starting** · 20 Dec 23, 18:40

Performing Pore Scan

**Pore scan result** · 20 Dec 23, 18:44

Pore scan for flow cell PAU00939 has found a total of 5064 pores. 1845 pores available for immediate sequencing

**Pore scan starting** · 20 Dec 23, 20:15

Performing Pore Scan

**Pore scan result** · 20 Dec 23, 20:19

Pore scan for flow cell PAU00939 has found a total of 4964 pores. 1830 pores available for immediate sequencing

**Pore scan starting** · 20 Dec 23, 21:50

Performing Pore Scan

**Pore scan result** · 20 Dec 23, 21:54

Pore scan for flow cell PAU00939 has found a total of 4731 pores. 1798 pores available for immediate sequencing

**Pore scan starting** · 20 Dec 23, 23:25

Performing Pore Scan

**Pore scan result** · 20 Dec 23, 23:29

Pore scan for flow cell PAU00939 has found a total of 4437 pores. 1759 pores available for immediate sequencing

**Pore scan starting** · 21 Dec 23, 01:00

Performing Pore Scan

**Pore scan result** · 21 Dec 23, 01:04

Pore scan for flow cell PAU00939 has found a total of 4174 pores. 1689 pores available for immediate sequencing

**Pore scan starting** · 21 Dec 23, 02:35

Performing Pore Scan

**Pore scan result** · 21 Dec 23, 02:39

Pore scan for flow cell PAU00939 has found a total of 3882 pores. 1581 pores available for immediate sequencing

**Pore scan starting** · 21 Dec 23, 04:10

Performing Pore Scan

**Pore scan result** · 21 Dec 23, 04:14

Pore scan for flow cell PAU00939 has found a total of 3610 pores. 1538 pores available for immediate sequencing

**Pore scan starting** · 21 Dec 23, 05:45

Performing Pore Scan

**Pore scan result** · 21 Dec 23, 05:49

Pore scan for flow cell PAU00939 has found a total of 3369 pores. 1482 pores available for immediate sequencing

**Pore scan starting** · 21 Dec 23, 07:20

Performing Pore Scan

**Pore scan result** · 21 Dec 23, 07:24

Pore scan for flow cell PAU00939 has found a total of 3181 pores. 1460 pores available for immediate sequencing

**Pore scan starting** · 21 Dec 23, 08:55

Performing Pore Scan

**Pore scan result** · 21 Dec 23, 08:59

Pore scan for flow cell PAU00939 has found a total of 2912 pores. 1493 pores available for immediate sequencing

**Pore scan starting** · 21 Dec 23, 10:30

Performing Pore Scan

**Pore scan result** · 21 Dec 23, 10:34

Pore scan for flow cell PAU00939 has found a total of 2686 pores. 1286 pores available for immediate sequencing

**Pore scan starting** · 21 Dec 23, 12:05

Performing Pore Scan

**Pore scan result** · 21 Dec 23, 12:09

Pore scan for flow cell PAU00939 has found a total of 2537 pores. 1167 pores available for immediate sequencing

**Pore scan starting** · 21 Dec 23, 13:41

Performing Pore Scan

**Pore scan result** · 21 Dec 23, 13:44

Pore scan for flow cell PAU00939 has found a total of 2395 pores. 1289 pores available for immediate sequencing

**Pore scan starting** · 21 Dec 23, 15:16

Performing Pore Scan

**Pore scan result** · 21 Dec 23, 15:19

Pore scan for flow cell PAU00939 has found a total of 2189 pores. 1031 pores available for immediate sequencing

**Pore scan starting** · 21 Dec 23, 16:51

Performing Pore Scan

**Pore scan result** · 21 Dec 23, 16:54

Pore scan for flow cell PAU00939 has found a total of 2049 pores. 979 pores available for immediate sequencing

**Pore scan starting** · 21 Dec 23, 18:26

Performing Pore Scan

**Pore scan result** · 21 Dec 23, 18:30

Pore scan for flow cell PAU00939 has found a total of 1935 pores. 1002 pores available for immediate sequencing

**Pore scan starting** · 21 Dec 23, 20:01

Performing Pore Scan

**Pore scan result** · 21 Dec 23, 20:05

Pore scan for flow cell PAU00939 has found a total of 1799 pores. 871 pores available for immediate sequencing

**Pore scan starting** · 21 Dec 23, 21:36

Performing Pore Scan

**Pore scan result** · 21 Dec 23, 21:40

Pore scan for flow cell PAU00939 has found a total of 1618 pores. 751 pores available for immediate sequencing

**Pore scan starting** · 21 Dec 23, 23:11

Performing Pore Scan

**Pore scan result** · 21 Dec 23, 23:15

Pore scan for flow cell PAU00939 has found a total of 1408 pores. 712 pores available for immediate sequencing

**Pore scan starting** · 22 Dec 23, 00:46

Performing Pore Scan

**Pore scan result** · 22 Dec 23, 00:50

Pore scan for flow cell PAU00939 has found a total of 1161 pores. 630 pores available for immediate sequencing

**Pore scan starting** · 22 Dec 23, 02:21

Performing Pore Scan

**Pore scan result** · 22 Dec 23, 02:25

Pore scan for flow cell PAU00939 has found a total of 907 pores. 498 pores available for immediate sequencing

**Pore scan starting** · 22 Dec 23, 03:56

Performing Pore Scan

**Pore scan result** · 22 Dec 23, 04:00

Pore scan for flow cell PAU00939 has found a total of 816 pores. 473 pores available for immediate sequencing

**Pore scan starting** · 22 Dec 23, 05:31

Performing Pore Scan

**Pore scan result** · 22 Dec 23, 05:35

Pore scan for flow cell PAU00939 has found a total of 727 pores. 422 pores available for immediate sequencing

**Pore scan starting** · 22 Dec 23, 07:06

Performing Pore Scan

**Pore scan result** · 22 Dec 23, 07:10

Pore scan for flow cell PAU00939 has found a total of 711 pores. 424 pores available for immediate sequencing

**Pore scan starting** · 22 Dec 23, 08:41

Performing Pore Scan

**Pore scan result** · 22 Dec 23, 08:45

Pore scan for flow cell PAU00939 has found a total of 961 pores. 639 pores available for immediate sequencing

**Pore scan starting** · 22 Dec 23, 10:15

Performing Pore Scan

**Pore scan result** · 22 Dec 23, 10:19

Pore scan for flow cell PAU00939 has found a total of 1138 pores. 743 pores available for immediate sequencing

**Pore scan starting** · 22 Dec 23, 11:49

Performing Pore Scan

**Pore scan result** · 22 Dec 23, 11:53

Pore scan for flow cell PAU00939 has found a total of 686 pores. 353 pores available for immediate sequencing

**Pore scan starting** · 22 Dec 23, 13:24

Performing Pore Scan

**Pore scan result** · 22 Dec 23, 13:28

Pore scan for flow cell PAU00939 has found a total of 526 pores. 245 pores available for immediate sequencing

**Pore scan starting** · 22 Dec 23, 14:59

Performing Pore Scan

**Pore scan result** · 22 Dec 23, 15:03

Pore scan for flow cell PAU00939 has found a total of 561 pores. 283 pores available for immediate sequencing

**Pore scan starting** · 22 Dec 23, 16:34

Performing Pore Scan

**Pore scan result** · 22 Dec 23, 16:38

Pore scan for flow cell PAU00939 has found a total of 631 pores. 349 pores available for immediate sequencing

**Pore scan starting** · 22 Dec 23, 18:09

Performing Pore Scan

**Pore scan result** · 22 Dec 23, 18:13

Pore scan for flow cell PAU00939 has found a total of 715 pores. 503 pores available for immediate sequencing

**Pore scan starting** · 22 Dec 23, 19:44

Performing Pore Scan

**Pore scan result** · 22 Dec 23, 19:47

Pore scan for flow cell PAU00939 has found a total of 568 pores. 288 pores available for immediate sequencing **UNIT ABBREVIATIONS**

Byte B Kilobyte KB Megabyte MB Gigabyte GB Terabyte TB

Base b Kilobase kb Megabase Mb Gigabase Gb Terabase Tb

Minutes mins Hours hrs

Generated using Run Report Template v.5.7.2
